# Evolutionary Insights Into Felidae Iris Color Through Ancestral State Reconstruction

**DOI:** 10.1101/2023.03.20.533527

**Authors:** Julius A. Tabin, Katherine A. Chiasson

## Abstract

There have been almost no studies with an evolutionary perspective on eye (iris) color, outside of humans and domesticated animals. Extant members of the family Felidae have a great interspecific and intraspecific diversity of eye colors, in stark contrast to their closest relatives, all of which have only brown eyes. This makes the felids a great model to investigate the evolution of eye color in natural populations. Through machine learning cluster image analysis of publicly available photographs of all felid species, as well as a number of subspecies, five felid eye colors were identified: brown, green, yellow, gray, and blue. Using phylogenetic comparative methods, the presence or absence of these colors was reconstructed on a phylogeny. Additionally, through a new color analysis method, the specific shades of the ancestors’ eyes were quantitatively reconstructed. The ancestral felid population was predicted to have brown-eyed individuals, as well as a novel evolution of gray-eyed individuals, the latter being a key innovation that allowed the rapid diversification of eye color seen in modern felids, including numerous gains and losses of different eye colors. It was also found that the gain of yellow eyes is highly associated with, and may be necessary for, the evolution of round pupils in felids, which may influence the shades present in the eyes in turn. Along with these important insights, the methods presented in this work are widely applicable and will facilitate future research into phylogenetic reconstruction of color beyond irises.

## Introduction

Eye (iris) color is one of the most conspicuous and varied traits among animals, at least for those that have colored irises. To date, much of the work investigating eye colors has focused on humans. This is not surprising, given how stark differences in human eye color can be, even between close relatives. This diversity of human eye colors, ranging from brown to green to blue, has been suggested to not be under strong natural selection (Jablonski and Chaplin, 2017). Thus, it has been attributed to bottlenecks and genetic drift, migration (Donnelly, *et al*. 2012), and especially sexual selection (Frost 2006). It is known that human eye colors differ due to a relatively small number of genes that act on the amount and quality of melanin in the eye (White and Rabago-Smith 2011; Simcoe *et al*. 2021). Yet, such eye color variation within a species (intraspecific variation) has been described as rare amongst animals, apart from artificially selected domesticated animals (Negro *et al*. 2017) and some species of birds. This latter variation has been mainly attributed to age, sex, or subspecies identity (Corbett *et al*. 2023).

In humans, the discrete view of intraspecific variation in eye color has also been challenged by more continuous models of variation (Liu *et al*. 2010). Traditional categories, such as blue, green, and brown, have been shown to overlap to varying degrees. This problem has been demonstrated for other color traits as well: for example, the colors of snail shells (Davison *et al*. 2019), tree lizard throats (Paterson and Blouin-Demers 2017), and bird plumage (Mould *et al*. 2023). These studies indicate that, although discrete categories are useful for trait comprehension and can facilitate clear analyses, results can be compromised if the categories do not truly reflect the real trait distribution. Some studies advocate for coupling analysis of discrete categories with more granular analysis within individuals of a category (Mould *et al*. 2023). Comprehensive investigation into the validity and utility of discrete categories for eye color has been lacking, especially since the optimal way to categorize color variation is still debated even for more well-studied traits (e.g. Cote *et al*. 2008; Vercken *et al*. 2008). In this way, not only has there been limited evidence for intraspecific variation in eye color beyond humans and domesticants, but it has also been unclear how to best categorize that variation, should it exist.

Even when just regarding eye color differences between species (interspecific variation), few hypotheses have been tested and very little is known about the adaptive benefits or evolutionary history of eye color, particularly in a natural context. While a few studies have attempted to tackle these questions by focusing primarily on distinctions between brightly or darkly colored irises, this method eschews quantitative color measurements and limits the conclusions that can be drawn (Craig and Hulley 2004). It is of particular interest to reconstruct the ancestral state of eye color because such reconstructions can shed light on the history of the trait and help interrogate how and why the current variants exist. Such analyses are vital for broadly increasing knowledge of evolution in a natural context, especially since eye color is not retained in fossils, nor in most preserved specimens. This has been done to great effect in owls; however, only “light” and “dark” eyes were considered in the analysis, not specific eye colors (Passarotto *et al*. 2018).

As with most groups, little work has been done understanding the eye colors of members of the family Felidae. Behavioral traits have been correlated to the eye colors of domesticated cats (*Felis catus*), but the wild felids have been largely left unstudied in this regard (Wilhelmy *et al*. 2016). This is surprising – although the closest relatives to the felids, such as linsangs, hyenas, and genets (Johnson *et al*. 2006), all have brown eyes and little inter- or intraspecific variation, the felids have a wide diversity of eye colors within and between them, even without counting *F. catus* (Figure 1). Although this trait is apparent by simply looking at members of each species, it has never been studied in an evolutionary context.

**Figure 1:**
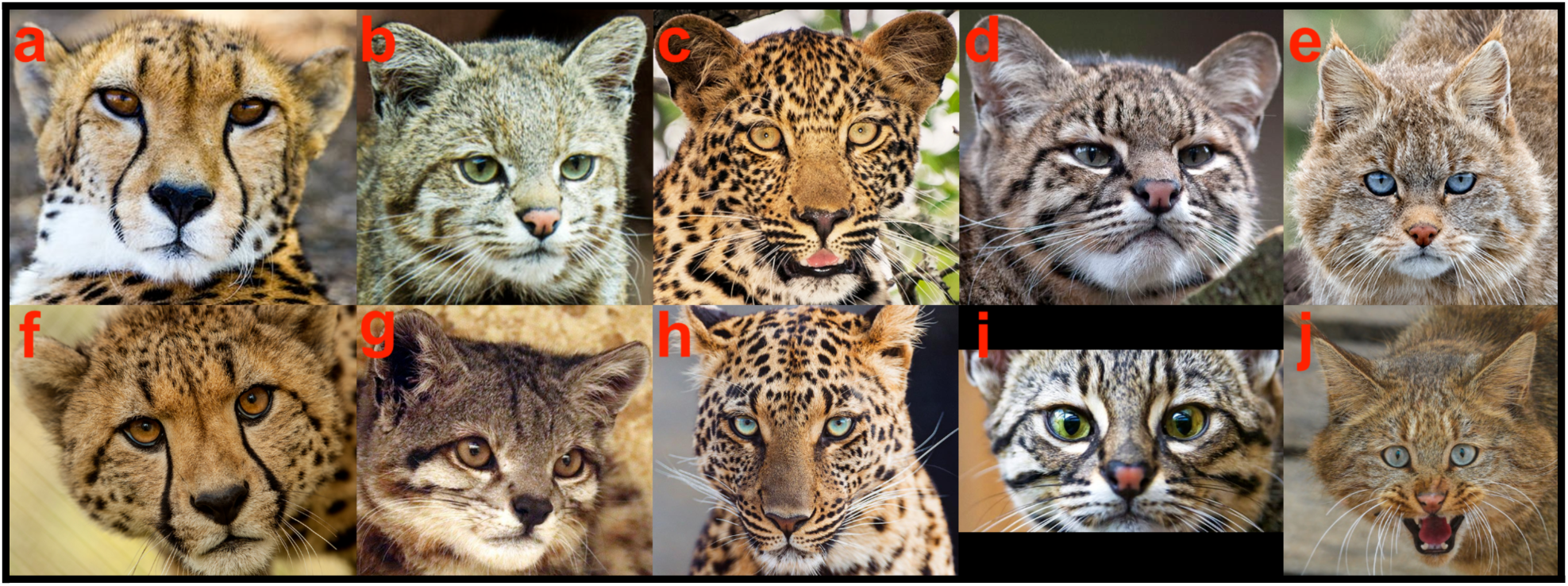
Examples of the five felid eye colors, with inter- and intraspecific variation. Each row contains an example of each of the five eye colors and each column contains two representatives of the same species. Colors and species are as follows: (a) brown (*Acinonyx jubatus*), (b) green (*Leopardus jacobita*), (c) yellow (*Panthera pardus*), (d) gray (*Leopardus geoffroyi*), (e) blue (*Felis bieti*), (f) yellow (*Acinonyx jubatus*), (g) brown (*Leopardus jacobita*), (h) blue (*Panthera pardus*), (i) green (*Leopardus geoffroyi*), (j) gray (*Felis bieti*). Photographs from (a) Piet Bakker, Pexels, (b) Luis D Romero, Shutterstock, (c) ABP News, (d) www.dinoanimals.com, (e) Song Dazhao, CFCA, (f) Ronda Gregorio, Smithsonian National Zoo, (g) Nayer Youakim, (h) Tambako The Jaguar, Flickr, (i) the Small Wild Cat Conservation Foundation, (j) Himimomi, www.zoochat.com.

Here, we present the first quantitative phylogenetic comparative analysis of eye color. We examine representatives from every extant felid species, as well as a number of subspecies, using a new quantitative color analysis method to solidify a categorization of eye color for these groups through discrete and continuous methods. Using this data, we have reconstructed the eye colors of the ancestors of the felids at all phylogenetic nodes, as well as tested their correlations with environmental data (zoogeographical region and habitat), behavioral data (activity mode), and morphological data (coat pattern, pupil shape, and pigmentation), to better understand the diversification of felid eye colors and demonstrate that phylogenetic investigations into eye color are not only possible, but fruitful.

## Results

To assess the range of eye colors present in the Felidae, we leveraged high-quality public image databases and sampled individuals from all non-domesticated felid species, as well as four related outgroups (see Methods for details). Which eye colors are present for each taxa was determined impartially using color identification software (Methods). Within the 52 felid taxa considered in the study, gray eyes were found to be present in 38 taxa (73%), brown eyes in 28 taxa (54%), yellow eyes in 23 taxa (44%), green eyes in 21 taxa (40%), and blue eyes in 6 taxa (12%). These statistics are given in Table 1, along with the results when subspecies are not considered and when just the most common eye colors (eye colors present in >80% of each taxon’s individuals in the surveyed databases; see Methods) are considered. Figure 1 contains representative images for each identified eye color.

**Table 1:** Eye color analysis count results

| Eye Color | In All Felid Taxa | Without Subspecies | Most Common Colors in All Felid Taxa | Most Common Colors Without Subspecies |
| --- | --- | --- | --- | --- |
| Gray | 38 (73%) | 31 (72%) | 31 (60%) | 23 (53%) |
| Brown | 28 (54%) | 28 (65%) | 22 (42%) | 22 (51%) |
| Yellow | 23 (44%) | 17 (40%) | 16 (31%) | 10 (23%) |
| Green | 21 (40%) | 20 (47%) | 11 (21%) | 10 (23%) |
| Blue | 6 (12%) | 5 (13%) | 2 (4%) | 2 (5%) |

In 10 felid taxa only a single eye color was observed, in 25 taxa two eye colors were observed, in 12 three eye colors were observed, and in 5 four eye colors were observed. All the eye colors present for each species are displayed in Figure 2. When considering just the most common eye colors in the populations of each taxon, 25 felid taxa had only a single eye color, 24 taxa had two eye colors, and 3 had three eye colors. Even with this conservative filtering of the data, there is conclusive evidence of the presence of intraspecific iris color variation among the Felidae.

**Figure 2:**
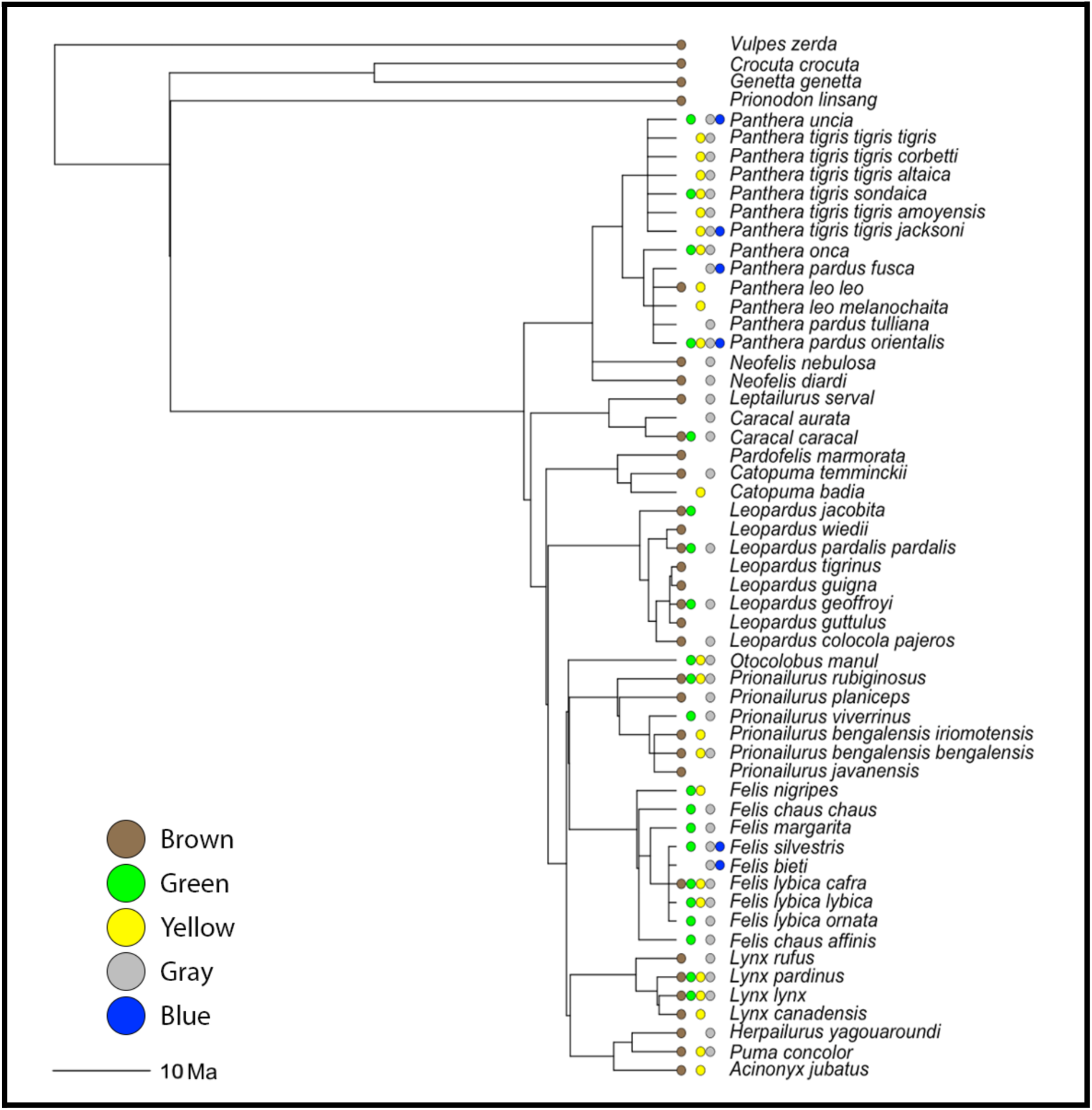
Phylogenetic tree of the Felidae (and outgroups), modified from Nyakatura and Bininda-Emonds (2012), with iris color data. Dots next to tips represent the presence of each eye color (black dot: brown eyes, green dot: green eyes, yellow dot: yellow eyes, gray dot: gray eyes, blue dot: blue eyes). No dot for a given color represents an absence of that color from the respective taxa. Graph created using plot.phylo() in ape.

The adequacy of analyzing felid eye color as a discrete polymorphic trait was supported by a principal component analysis on the RGB values of all the pixels in the data set (Methods). Principal component (PC) 1 was found to separate dark from light pixel coloration (Figure S1) and only brown eyes were found to be a meaningfully distinct category along that axis (p<0.001, Satterthwaite’s t-test on a linear mixed model with Bonferroni correction; Supplementary Table 1). In other words, the data set photos of brown eyes contained significantly darker pixels than the other colors, unsurprising given their darker pigment. While pigment darkness is a biologically relevant metric for eyes (Passarotto *et al*. 2018), this axis is not sufficient to judge the biological validity of discrete color categories since it only encapsulates a singular aspect of color variation. Two eyes with unambiguously distinct colors could still have the same darkness value. An axis that directly contains the colors of interest is far more relevant to determine whether the categories significantly separate: if one wants to evaluate whether blue and gray, for instance, are different categories in the data set, the axis of measurement must thus directly contain blue and gray.

PC2 was found to separate blue, gray, brown, and yellow pixels and all color categories were significant along that axis (p<0.0001 for all groups, as well as for all group comparisons, Satterthwaite’s t-test on a linear mixed model with Bonferroni correction and a post-hoc Tukey HSD test with Bonferroni correction, respectively; Supplementary Table 1). There was little overlap between the pixels within the categories along PC2, except for green, which overlaps with brown and gray. However, PC3 was found to separate green pixels from other pixel colors. In this case, only green eyes were found to be meaningfully different along the axis (p<0.001, Satterthwaite’s t-test on a linear mixed model with Bonferroni correction). In sum, the first three PCs comprised 100% of the variance in pixel coloration. Thus, for the data set as a whole, the presence of the five discrete color groups was supported and those color groups were able to be easily distinguished along relevant axes. Support for the discrete color categories was also found for every species in the data set when analyzed individually along PC 2 (Methods), except for *Acinonyx jubatus* (p=1.00, Kruskal–Wallis test with Bonferroni correction; Figure S2; Supplementary Table 1). Images of color variation in *Acinonyx jubatus* can be seen in Figure 1a and f.

### General color reconstruction

Eye color was reconstructed by subsetting a published ultrametric Carnivora supertree (Nyakatura and Bininda-Emonds 2012; see Methods for details). This “main phylogeny” is nevertheless missing nine extant felid taxa, which were added manually (Methods) to generate an expanded “full phylogeny”. The overall ancestral state reconstruction for all of the colors on the main phylogeny is given in Figure 3, with a color considered present if there was greater than 50% support as determined by maximum likelihood methods (i.e. more likely than not). This reconstruction does not substantially differ from the reconstruction for the full phylogeny, with additional species and subspecies added (Figure S3). The only differences are that, for the full phylogeny, there are fewer blue-eyed ancestors on the tree and there is less confidence that the common ancestor of the Ocelot lineage had gray eyes. Most of the presence/absence information for the colors have high maximum likelihood support across the tree (Figure S4). However, a notable exception is green. Under the main tree, taking into account all observed eye colors, the confidence in the presence of green is around 50% for every node, likely due to the wide and seemingly unpatterned distribution of green eyes across taxa. This lack of confidence in the presence of green eyes is ameliorated when either using only the most common eye colors or the full phylogeny, but not when alternate models of trait evolution were tried. In both cases, the models predict the presence of green eyes in the ancestor of the Domestic Cat lineage with high confidence. The analysis using only the most common eye colors predicts green eyes going back three more common ancestors. Apart from these few areas, all three analyses are congruent. The reconstruction for all five colors are consistently unclear for the ancestor of the Felidae and *Prionodon linsang*, as well as any of the deeper nodes on the tree. This is unsurprising, given that ancestral state reconstruction becomes more uncertain the farther back one goes.

**Figure 3:**
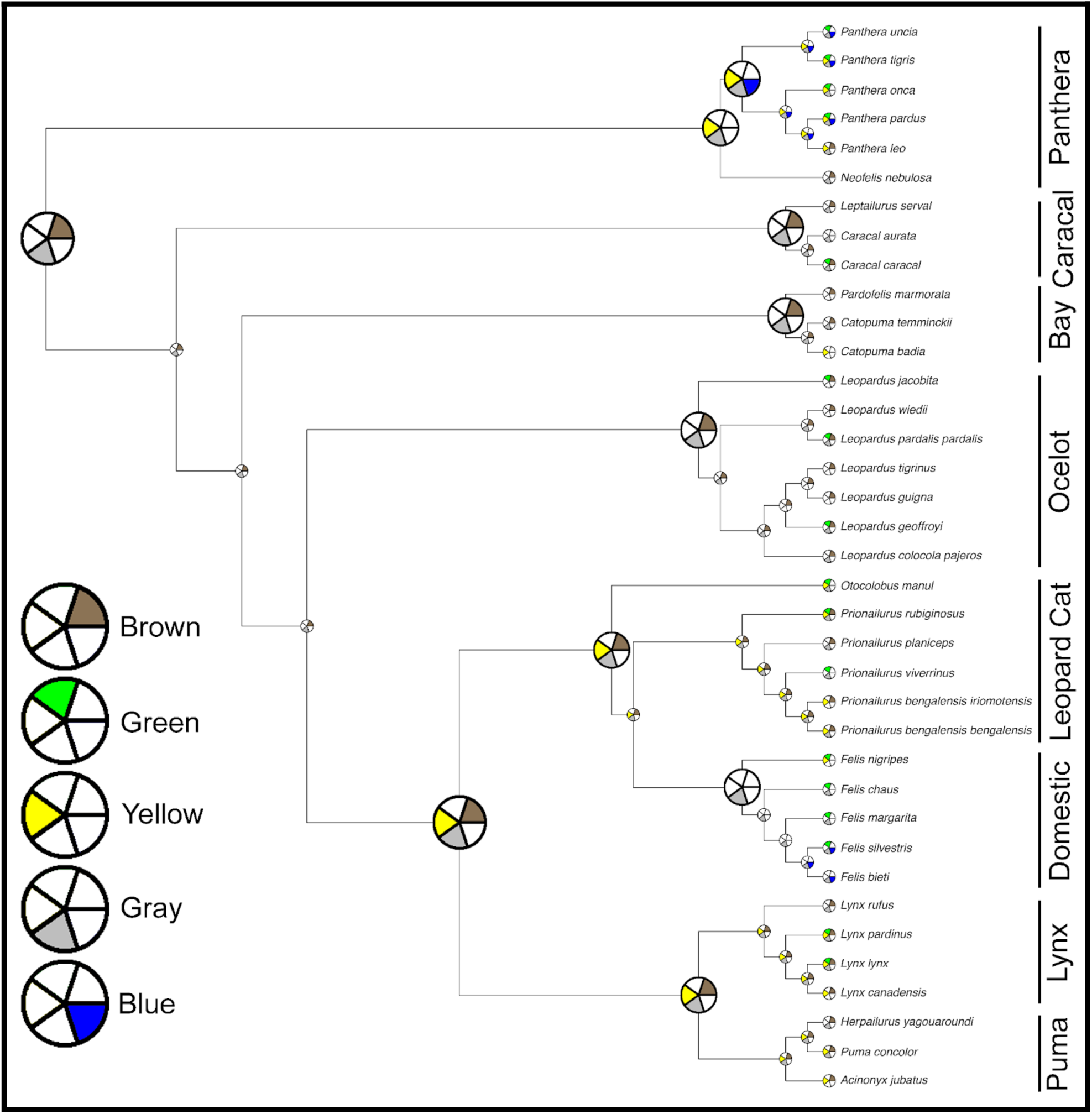
Reconstruction of the ancestral states of all five eye colors for the Felidae. The five-wedge pie charts indicate presence (color) or absence (white) of the various iris colors. Colors were considered present if there was >50% support as determined by maximum likelihood methods (probabilities plotted in Figure S4). Exact branch lengths are not plotted and lineage names are given on the right.

The ancestor of the Felidae is reconstructed with high likelihood to have had both brown- and gray-eyed individuals present in its population. This represents two novel changes: having multiple eye colors in the same species (intraspecific eye color variation) and having gray eyes in particular. There is good evidence that this is the only major gain of the gray-eyed trait in the Felidae, although a few taxa subsequently lost it. Brown eyes are also common throughout the tree, but had two major losses, once when the Domestic Cat Lineage diverged 6.2 million years ago and once after the Panthera Lineage diverged 10.8 million years ago (Johnson *et al*. 2006). However, there is uncertainty and discordance between the three analyses about whether the Panthera Lineage ancestor had brown eyes, so this latter brown eye loss may have occurred after the genus *Panthera* split from *Neofelis* approximately 6.5 million years ago. As with gray eyes, there were multiple other species-specific losses, as well as a regaining of brown eyes by *Panthera leo*.

The presence of yellow eyes is predicted to have convergently evolved at least twice in ancestors and multiple more times in individual extant species. The two demonstrated higher than species-level gains of yellow eyes are when the Panthera Lineage split off and when the Lynx, Leopard Cat, Domestic, and Puma Lineages diverged from the rest of the tree (8 million years ago) (Johnson *et al*. 2006). Much like the other colors, there were a number of cases of loss and even regaining of yellow eyes.

The presence of green eyes stands apart from the previous three colors because it does not seem to have developed early in the evolution and diversification of felids. Instead, it appears to have evolved at least twelve individual times, most of the time at the species level alone. According to the congruent part of the most common color and full phylogeny analyses, the most significant development of green eyes occurred in the Domestic Cat Lineage when it diverged, only being lost once in that lineage (*Felis bieti*). In fact, that would be the only observed time green eyes were ever lost in the Felidae. The presence of blue eyes has a similar evolutionary distribution to that of green eyes, albeit much more rare, possibly having evolved independently at least twice, according to the main analysis: in the ancestor of the *Panthera* genus and in the ancestor of *Felis silvestris* and *Felis bieti* (also the ancestor of *Felis catus*, not considered in the phylogeny). It should be noted that the most common color and full phylogeny analyses both lead to an alternative prediction that the blue eye color was not present in any of the ancestors and arose in individual tips independently.

### Quantitative color reconstruction

Beyond assigning eye color to one of the aforementioned five broad color groupings, RGB values from each image were processed using a dimensionality reduction algorithm and examined using a cluster analysis (see Methods for details). This is needed because, while the broad color categories are distinct, it is rare that any given felid eye is homogenous in color (see Figure 1b, c, and h for clear examples). This analysis resulted in a quantitative and finely detailed output of the average number of shades for each eye color in each taxon, as well as what the colors of those shades are. All eye colors for all species had between two and four shades (see Figure 1f and 1h for examples of eyes with two and four shades, respectively). Our stringent image selection criteria was shown to provide accurate and robust assessment of shade coloration, even at very low sample sizes, with little variation due to lighting (Figure S5).

The reconstruction of the shades of brown eyes, conducted with reference to the presence/absence reconstruction done above, reveals some large-scale evolutionary trends. The eye color of the outgroups (*Vulpes zerda, Crocuta crocuta*, *Genetta genetta*, and *Prionodon linsang*), as well as their close non-felid relatives that were not analyzed in this study, strongly suggests that the ancestor of these groups likely had particularly dark brown eyes, as it is unlikely that similar dark brown eyes would have convergently evolved across closely related and ecologically distinct species. Our study does not have the requisite power to confirm the presence of brown eyes for the common ancestor of the whole tree, but our models do predict that the subsequent ancestors of the outgroups and the Felidae had brown eyes. The predicted darkness is also recapitulated in the reconstruction, including for the common ancestor of the whole tree, if it is assumed to have had brown eyes (Figure 4a). The brown eye colors of all the outgroup ancestors are reconstructed as quite dark, albeit not as dark as the irises of *Prionodon linsang*. The brown eyes of the ancestor of the felids are predicted to have had a lighter coloration, but still with the same shade distribution as the deeper ancestors: the medium shade is the primary shade in the eye, followed by light and dark. After this, the proportion of light, medium, and dark shades changes frequently in the tree. In the data, the medium shade nevertheless remains most likely to be the primary shade (5 for light, 20 for medium, and 7 for dark). The three shades have about an equal distribution of being the secondary shade (10 for light, 11 for medium, and 11 for dark). The overall shade of brown eyes (taking into account dark, medium, and light shades for each) also undergoes substantial changes over the tree. In some lineages, such as the Lynx or Ocelot Lineages, the shade returns to a darker state, as it was before felids branched off. In other lineages, such as the Leopard Cat Lineage, the shade continued to lighten.

**Figure 4:**
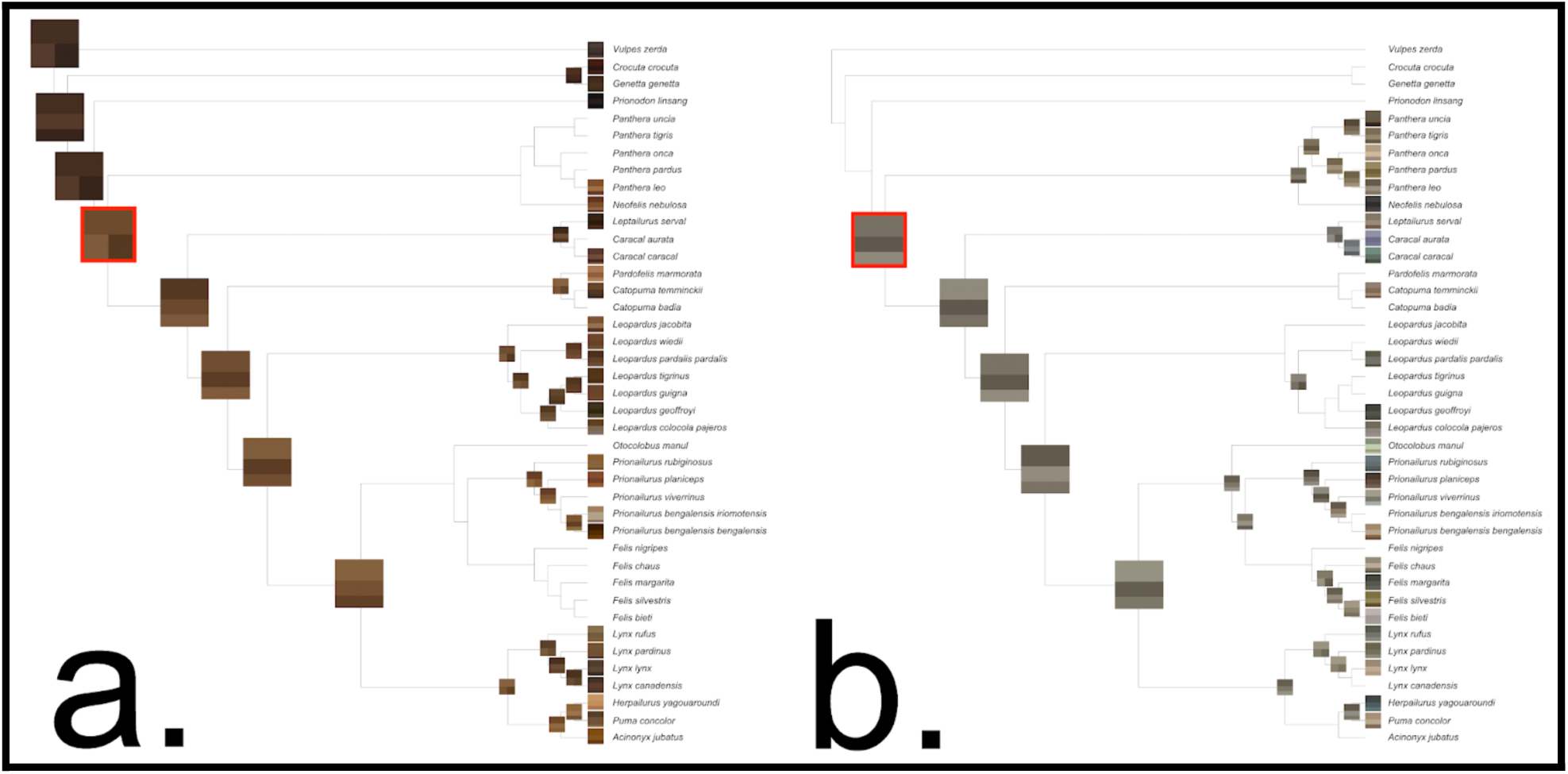
Reconstruction of the ancestral states of the shades of (a) brown and (b) gray eyes. The squares at each node are the quantitative reconstructed shades. The proportion of a square that a shade takes up indicates how common that shade is in the data. Exact branch lengths are not plotted. The square highlighted in red is the ancestor of the Felidae.

A high level of variation in gray eyes is apparent from viewing the types of gray in the data (Figure 4b; Supplementary Results). Unlike brown, where all of the variety was focused within a relatively narrow region, there are gray colors that span a large spectrum, being closer to brown, green, blue, or yellow for different taxa. The gray that was reconstructed for the ancestor of the Felidae (RGB: 118, 112, 100) is closer to brown-gray than pure gray, a trait that continues as the Felidae diversified. Gray colors have close to equal red, green, and blue values, whereas brown colors have much higher red and green values than blue. The ancestral gray having a brownish character is evident by the decreased blue value, compared to the red and green values. The brown content in gray-eyed animals is particularly strong in the Panthera Lineage, eventually nearly becoming fully brown for certain taxa. In the Domestic Cat Lineage, the gray color substantially lightened, losing its brown content and sometimes taking on a slightly higher green content, particularly for *Felis silvestris*. When the genus *Caracal* split from the rest of the Caracal Lineage, its gray changed to have much higher blue and green content (for *Caracal aurata* and *Caracal caracal*, respectively). More blue content in the color of gray eyes is a repeated trait, occurring for *Prionailurus rubiginosus* and *Herpailurus yagouaroundi* as well. For gray, the dark shade is most commonly the primary shade (15 times), closely followed by medium (14 times), then light (9 times). This order is almost reversed for the secondary shades, although it is closer (14, 16, and 8 for light, medium, and dark). Similar analyses of the evolution of yellow, green, and blue eyes are shown in the Supplementary Results, as well as Figures S6-8.

### Correlation analysis

The presence of the five eye colors were correlated against one another, taking the phylogeny into account (Figure 5a). The correlations for the main phylogeny with all observed eye colors sometimes differ from the results with only the most commonly observed eye colors (Figure S9a) or for the full phylogeny (Figure S9b), but often there is agreement. For all three analyses, a significant positive correlation (log Bayes Factor > 2) was identified every time a color was correlated with itself, a positive baseline check of the quality of the method. The main analysis demonstrated a significant negative correlation between the presence of brown eyes and the presence of both green (BF = 8.33, corr = -0.71) and blue eyes (BF = 15.38, corr = -1; indeed, no felid species has both brown and blue eyes). In contrast, the presence of gray eyes is significantly positively correlated with the presence of both green (BF = 7.04, corr = 0.63) and blue eyes (BF = 5.26, corr = 0.98; all blue-eyed taxa also have gray eyes). These correlations for gray, as well as the negative brown-blue correlation, were found for the other two analysis types as well. However, the analysis with just the most common eye colors indicated additional negative correlations between brown eyes and both yellow (BF = 8.87, corr = -0.78) and gray eyes (BF = 7.01, corr = -0.67).

**Figure 5:**
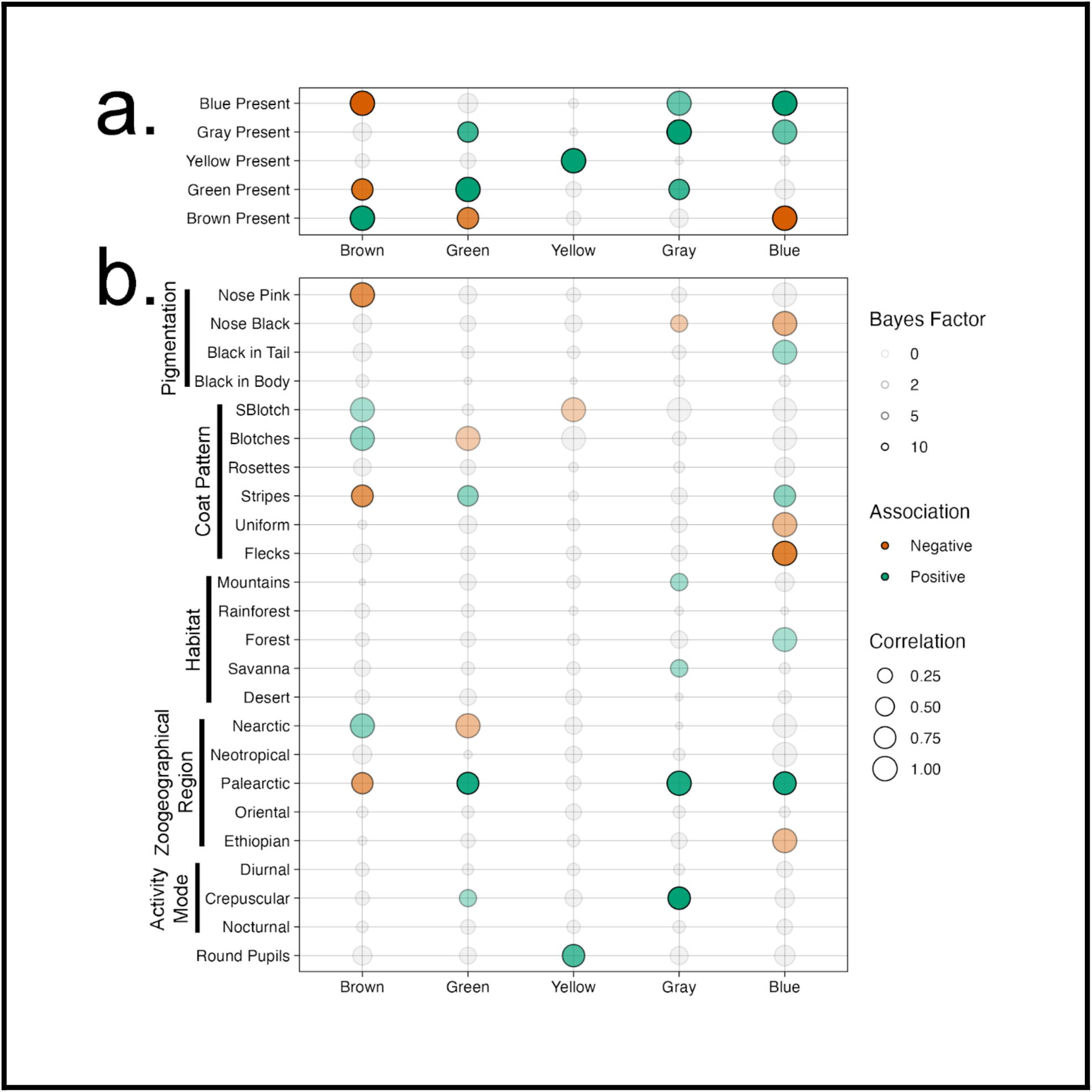
Correlations between the (a) presence of each eye color and (b) against various physical, behavioral, and environmental factors. Larger circles correspond to stronger correlations and more opaque circles correspond to more significant correlations. Green circles have a positive correlation, red circles have a negative correlation, and gray circles do not meet the significance threshold (Bayes factor = 2).

Correlation analysis was also carried out on eye color relative to a variety of other physical traits (e.g. pupil shape, coat color), behavioral traits (e.g. diurnal versus nocturnal activity), and habitat characteristics (Figure 5b). Overall, most environmental and physical factors considered in the analysis showed at least some significant correlations with various eye colors, indicative of the complexity of eye color evolution. Notably, none of the activity modes were correlated with any eye color, except for crepuscularity with gray and green eyes—good evidence that this trait is not particularly important for eye color evolution in felids. Additionally, of the significant correlations, gray eyes were almost always positively associated with other traits. This greatly contrasts with the other eye colors, all of which have closer to a 50/50 distribution of positive and negative associations.

When eye colors were correlated to pupil shape, one significant correlation appeared: the presence of yellow eyes is strongly positively correlated with round pupils (BF = 6.32, corr = 0.80). This correlation was found in both other analyses as well. For all felid taxa that have round pupils, yellow eyes had evolved prior and, even if the taxa subsequently lost yellow eyes, yellow eyes are predicted to have been present in all ancestors that evolved round pupils. Given the above potential negative relationship between brown and yellow eyes, it is also unsurprising that only two taxa evolved round pupils while already having brown eyes (the closely related *Acinonyx jubatus* and *Puma concolor*). It should be noted that one can flip the direction of these correlations to obtain the associations with vertical pupils, given that this is the only other pupil type for felids.

There were some correlations between zoogeographical regions and various eye colors that cannot be otherwise explained by phylogeny, particularly on the ends of the color spectrum: brown eyes are positively associated with the nearctic region (BF = 3.77, corr = 0.97), gray eyes are positively associated with the palearctic region (BF = 8.39, corr = 0.99), and blue eyes are negatively associated with the neotropical region, albeit only in the two alternate analyses (BF = 4.02/4.70, corr = -0.95/-0.96). On the other hand, there were few correlations with habitat. For more on these results, including the shade correlations given in Figure S10, see the Supplementary Results.

## Discussion

Our image-based analysis demonstrates that the family Felidae has a broad range of interspecific and intraspecific variation in eye color, the latter type of variation being conspicuously absent in all the group’s close relatives. Indeed, within-species eye color variation has been thought to be largely absent from wild animals as a whole, except for those species where eye color varies on specific axes, such as age or sex (Negro *et al*. 2017). While this rule may hold for most groups, the Felidae constitute a notable exception, with over 80% of the taxa surveyed in this study having two or more different eye colors in their populations. The images that make up the data set were controlled to be all adults, so this cannot be due to a difference between juvenile and adult morphs. Although the sex of animals in the data set could not be determined, sex alone also cannot account for the variation seen, since 33% of taxa had more than two eye colors in their population. Variation in eye colors to this extent has not been formally described, except in humans and domesticated animals, making the felid system an ideal model to investigate the evolution of eye color.

Our results support the existence of discrete, polymorphic eye color categories for felids, with pixels from brown, green, yellow, gray, and blue eyes separating along biologically meaningful axes, with little overlap between these categories overall. This separation was present for the data set as a whole, as well as for every species but one. The outlier is *Acinonyx jubatus*, for which there was no statistical support for a distinction between brown and yellow eyes. The fact that this unique overlap between the “yellow” and “brown” categories for *A. jubatus* is not seen in its closest relatives, such as *Puma concolor*, likely indicates that these color categories have recently become more similar in this species. This may be attributed to the lack of genetic diversity in the species and the high level of inbreeding, leading to variation in color being difficult to maintain (Merola 1994).

Despite felid eye color being broadly discrete, pigment levels can nevertheless be continuous and prior studies on color variation suggest that even traits that can be discretized may have continuous variation at a more fine-grain level (Mould *et al*. 2023). Our finding that there are always at least two shades present within each eye supports this, providing a useful and biologically realistic model for future research into the evolution of eye color variation: treating the overall trait as fully discrete and then treating each category as continuous. This avoids unrealistic transitions between colors (see Methods for an example) in the overall model, while still allowing within-color variation to be analyzed. The quantitative shade analysis also allows eyes with substantial color differences within them to be separated (for examples, see the eyes of *Panthera pardus* in Figure 1c and h). This variation is retained in our continuous, quantitative analyses (see *P. pardus* in Figure S8). If eye color was treated only as a discrete category, these differences would be ignored, a particularly sizable loss if an eye’s coloration includes colors contrary to the overall color group assigned based on the majority of the eye (for more discussion about such eyes, see Supplementary Results).

The reconstruction of eye color indicates with high likelihood that the common ancestor population for the Felidae had both brown- and gray-eyed individuals. The presence of brown eyes is not surprising, given that all close relatives of the Felidae have dark brown eyes with no intraspecific variation. However, the presence of gray eyes is likely a family-specific characteristic. Although our analysis has left it unclear whether the ancestor of the Felidae and *Prionodon linsang* included gray eyed individuals, given that all of the close relatives of the Felidae (e.g. genets, hyenas, etc.) have only brown eyes, it is highly likely that this uncertain ancestor had exclusively brown eyes as well (Johnson *et al*. 2006).

The gray eye color is likely an intermediate between all of the other eye colors. Eye color is determined by the amounts of the pigments eumelanin and pheomelanin in the iris (Kolb *et al*. 2011). In a simple view, eyes with more eumelanin are brown, eyes with more pheomelanin are yellow, and eyes with lower levels of pheomelanin and eumelanin are blue or green. Gray eyes contain a moderate amount of both pigments, but not enough of either one to reliably be placed in another color group. This is supported by gray eyes in the data having much higher variability than the other four colors. If a population, such as the relatives of the felids, is homogenous for dark brown eyes, thus having a high level of eumelanin and little pheomelanin, it would be difficult to suddenly develop blue eyes, given that blue eyes need a very specific balance between the two pigments that is far from the dark brown state. Even a total loss of pigment, as with albinism, could not account for this, because a certain amount of pigment is still needed to have the blue color be visible (White and Rabago-Smith 2011).

Under this view, once gray eyes evolved in the felid ancestor, it became far easier to transition between eye colors and evolve new ones, resulting in the great diversification seen in the Felidae. It is out of the scope of this study to answer exactly which genetic changes led to this, but this is a question that should prompt future research. A promising starting point is identifying and comparing orthologous sequence data for genes known to affect melanin production in other species, such as OCA2, HERC2, and MC1R, in as many felid species as possible, to attempt to pinpoint felid-specific genetic changes that might affect eye colors (White and Rabago-Smith 2011).

Further evidence for the evolution of gray eyes being an intermediate form, stemming from a fully brown-eyed population, can be found in the shade reconstruction. The ancestral felid’s gray eyes were not purely gray but were made up of brownish-gray shades. This is only plausible if there is a gradient from brown to gray with no other colors in between and if the gray eyes were formed from a modified brown eye. Furthermore, when examining the quantitative shades of gray across the phylogeny, there are places where other colors were lost, coupled with a shift towards that color by gray. For example, in the genus *Panthera*, when brown was lost, there was a concurrent increase in the amount of brown in the gray-eyed animals. This could be explained by a gradual change of brown eyes becoming more gray, merging the two colors, or by discrete changes, with brown eyes being lost and then gray eyes becoming browner. By the present day, the gray eyes in the *Panthera* have almost crossed back into being brown (for example, *Panthera tigris*). Additionally, there are a number of species for which the content of blue has substantially increased in their gray eyes, such as for *Herpailurus yagouaroundi*, *Prionailurus rubiginosus*, and both species in the genus *Caracal*. However, this is never the case for taxa that already have blue eyes, all of which also have gray eyes.

In contrast to *Panthera*, when the Domestic Cat Lineage lost brown eyes, it was coupled with a lightening of the color of gray eyes and likely the evolution of green eyes. In this case, neither of the present colors are close to brown. This represents a second path for the loss of brown eyes: rather than occupying the place of brown eyes in the population by effectively merging brown and gray, the population of the ancestor of the Domestic Cat Lineage shifted the entire eye color scheme. This likely requires a greater number of changes and it is no wonder that such examples of huge eye color scheme shifts are rare. Through comparisons of this nature, the data collected and analyzed in this study can provide important insights into eye color evolution on both small and large scales. It should also be noted that many of the wild species within the Domestic Cat Lineage can breed with the domestic cat (*Felis catus*) (Oliveira *et al*. 2008; Lyons 2012). Albeit unlikely, disruption from this form of hybridization is possible, given that many domestic cats have artificially selected eye colors.

Although our method of placing eyes into discrete color categories and then subsequently categorizing their variation in a continuous way is supported by the data, the quantitative reconstruction is subject to a relevant limitation. Since ancestral state reconstruction is applied to each color separately, species with eyes near the border between two color categories are not taken into account in both reconstructions. For example, the ancestor of the Panthera Lineage is predicted to have lost brown eyes and concurrently gained yellow eyes. If yellow eyes directly and gradually evolved from the brown eyes of the ancestor’s ancestor and the new yellow is simply a different shade of brown, then the shades of the yellow eyes of the Panthera Lineage should be taken into account when reconstructing the brown shade of the ancestor’s ancestor. Although this is an unlikely scenario, it is worth noting that it is beyond the scope of our method to account for it unless the discrete categories are ignored, which would lead to a host of far more pervasive and consequential issues.

The correlation results were also revealing. It is unsurprising that brown and blue eyes, at nearly opposite ends of the pigment spectrum, do not frequently coexist in natural populations and are significantly negatively associated. On the other hand, gray eyes, being an intermediate which is bordering the blue color space, provide an ideal anchor for the rarer blue eyes. The maintenance of blue eyes would be much more likely if blue-eyed individuals mated with blue-eyed or gray-eyed individuals, rather than with brown-eyed individuals. No felids naturally have both brown and blue eyes, so a worthwhile future direction would be to empirically test whether there is segregation in mating preference along eye color lines. Mating preferences provide an intriguing possibility for the evolution of eye color differences, given that one hypothesis for the diversity of human eye colors is sexual selection (Frost 2006). Cats are dichromatic and cannot recognize reds and oranges, but they can distinguish other colors (Clark and Clark 2016). This range of color sensitivity fits well, given that all the eye colors identified in this study are likely visible to felids and would thus be possible to be sexually selected for. However, even if this was the driving factor behind eye color diversification, it still does not explain the emergence of gray eyes, nor the differences between lineages.

Yellow eyes are also much more likely to coexist with round pupils. Round pupils are a repeated innovation in felids (the ancestral felid had vertical pupils; Banks *et al*. 2015), so it is notable that they only appeared in ancestors that are predicted to have already developed yellow eyes. Thus, the gain of yellow eyes acts as a prerequisite for the evolution of round pupils. However, our analysis cannot distinguish between correlation and causation. It seems probable that either an aspect of yellow eyes (causation), or an undiscovered lifestyle or physical trait that strongly covaries with yellow eyes (correlation), aligns with the conditions that are ideal for round pupil evolution. While not every species with yellow eyes developed round pupils, the two traits are tightly linked and only *Panthera uncia* lost yellow eyes after evolving round pupils. The reverse of this trend seems to be true for brown eyes, since only two groups had brown eyes before evolving round pupils. This makes sense, given the negative association between brown eyes and yellow eyes in the analysis with only the most common eye colors.

The contrasting forces of brown and yellow eyes can be seen in the shade correlations, with yellow eyes lightening the overall red and green shades of a species and brown eyes darkening the overall red, green, and blue shades. If gray eyes developed by decreasing the amount of eumelanin in the eyes, it could be that a second change increased pheomelanin levels, leading to yellow eyes. Then, with a new type of darker pigmented eye, there might have been less of an evolutionary “need” for eyes with lots of eumalanin. This could explain why many species have either brown or yellow eyes, but not both, particularly in lineages that simultaneously gained yellow and lost brown, such as the genus *Panthera*. For such lineages, the concurrent loss of brown and gain of yellow could also have been achieved if the second change altered the melanin synthesis pathway in brown eyes (as opposed to yellow stemming from gray). This could be achieved by increasing the likelihood that their shared precursor, dopaquinone, becomes pheomelanin, rather than eumelanin (Ito *et al*. 2000). Either way, the divergent effects of yellow and brown on round pupil evolution fit with these two colors being unlikely to develop together, but not being mutually exclusive.

There were few significant correlations for activity modes, even with the shade data taken into account. This is surprising, given the findings of Passarotto *et al*. (2018), which found that darker colored eyes in owls evolved in response to the switch to a nocturnal lifestyle. This is clearly not the case for felids. The ancestral state for felids is nocturnality, but gray eyes (usually lighter than brown eyes) evolved before any taxa made the switch to diurnality (Myers *et al*. 2022).

The fact that there were a number of significant correlations by zoogeographical region is fascinating, given how large each region is. This, coupled with the lack of significant correlations found for most habitats and the uniformity of eye colors across most animals around the world, indicates that the physical environment may play less of a direct role on eye color in felids and possibly mammals as a whole. Future investigations should be done to map eye colors at a population level. This mode of data collection, ideally paired with genomic analyses, would enable full phylogeographic investigations into the evolutionary history of the trait, beyond the broad correlations presented here.

Other traits, such as social system and mode of hunting, are also uninformative, given that the only long-distance (as opposed to short-range ambush) predator felid is the cheetah and the only non-solitary felid is the lion, neither of which show unique eye color characteristics (Banks *et al*. 2015; Myers *et al*. 2022). Thus, the specific adaptive benefit of having different eye colors is left as an open question.

It is known that eye color is at least partly tied to coat color in domestic cats and some similar associations do appear in our data, such as having brown eyes being negatively correlated with having a pink nose (Strain 2007). Having a pink nose, an easily measurable partial stand-in for a de-melanated skin color, is unsurprisingly not frequently found in species with brown eyes, which require more melanin. The same logic explains the negative correlation between blue eyes (which need lower melanin) and a black nose (which needs higher melanin). However, for the most part, the color or shade of felid eyes is not related to skin or fur color. This lack of coupling of the two traits, apart from the most and least melanated cases, likely allowed for the independent evolution of gray eyes in the felid ancestor.

All of the evidence presented here supports a larger theory of felid eye color evolution. Through random, novel mutation(s) that decreased the levels of iris eumalanin, a subset of the population of the ancestor of the Felidae developed gray eyes. Once this key innovation occurred, the new standing variation led to rapid diversification. Yellow, green, and blue eye colors evolved repeatedly, albeit not necessarily through the same genetic changes, as felid lineages diverged and groups reached new zoogeographical regions. The presence of these colors and the strength of the shades within them varied tremendously through interactions with different environments and physical characteristics. Tradeoffs between the amounts of pigment in the iris created antagonistic relationships between blue and brown eyes, as well as yellow and brown eyes, making their coexistence in various species less likely. The yellow-brown tradeoff, influenced by a potential increase in pheomelanin, possibly affected the development of round pupils. Iris color diversification represents a defining feature of the Felidae family and the data presented here demonstrates the complexity of the trait.

Eyes, and especially eye colors, have historically been a missed opportunity for evolutionary research. Because they are a rarely preserved element of animal bodies, some have assumed that eye color diversification over time can only be studied through ancient DNA (Negro *et al*. 2017). In contrast to this view, the work presented here demonstrates that eye color can be studied in an evolutionary context, without the need for ancient DNA or even DNA information at all. Through this work, the evolution of eye colors in the Felidae is now much clearer and there are many avenues for more studies, particularly regarding the clearly important evolutionary place that gray eyes occupy. This study provides a starting point for future research into eye color evolution in natural populations, a question that has not had any significant investigations until now, and could be easily expanded to other taxa. In addition, the scope of this study could be built upon by adding genetic data to the correlation analysis to work towards answering more functional questions. Beyond this, the method for quantitative color reconstruction presented in this study could be adapted to any color-based analysis, even beyond the iris. This will allow for high precision color reconstructions that were previously impossible.

## Limitations of the Study

While ancestral state reconstruction can provide statistical insight into the past, it remains a prediction. This study does not conclusively prove that the ancestor of the Felidae had gray eyes; it strongly predicts that this was the case, given modern data. For taxa where ancient DNA is available from an ancestral population, such as for humans, predictive models can be used to robustly predict eye color information (e.g. Draus-Barini *et al*. 2013). Thus, one way to be more sure about the eye colors of felid ancestors would be to use the phylogenetic comparative methods presented here in concert with ancient DNA, allowing the predictions made by the models used here to be empirically tested. Of course, a prerequisite to this study would be gaining a firm knowledge of which genes influence modern felid eye color and how their pathways interact, as well as extensive ancient DNA sampling of extinct felids.

Additionally, despite online databases providing essential support for this work, using such databases limits the amount of data that can be collected on possible confounders. For instance, the age, sex, and provenance of the animals whose photos were used for this study could not be reliably determined. This is especially relevant for rarer species, which are underrepresented in the data set for this study. While these factors should not have affected the overall conclusions of this study, data was not able to be collected about how eye color varies on these biologically-relevant axes.

Lastly, the sampling done for this study is only sufficient to make robust predictions about the Felidae. For a researcher interested in species beyond this family, more data would have to be collected and analyzed using the methods described in this article.

## Supporting information

Supplemental Results

## Acknowledgements

We would like to thank Clifford Tabin for methodological and manuscript preparation assistance. We want to thank Scott Edwards and his OEB 275R Comparative Genomics seminar, Landen Gozashti, Carolyn Elya, and the members of the Hoekstra Lab at Harvard University for conceptual and methodological assistance. We also would like to thank Liam Revell, Jeremy Beaulieu, Jonathan Schmitt, and especially Michaël Nicolaï for their useful advice on refinements of the methodology.

This work was supported in part by a graduate stipend from the Department of Organismic and Evolutionary Biology at Harvard University.

## Author Contributions

Conceptualization, J.A.T.; data collection and preparation, J.A.T. and K.A.C.; data analysis, J.A.T.; writing – original draft, J.A.T.; writing – review & editing, J.A.T. and K.A.C.; supervision, J.A.T.

## Methods

### Data set

In order to sample all felid species, we took advantage of public databases. Images of individuals from 40 extant felid species (all but *Felis catus*, excluded due to the artificial selection on eye color in domesticated cats by humans), as well as 12 identifiable subspecies and four outgroups (banded linsang, *Prionodon linsang*; spotted hyena, *Crocuta crocuta*; common genet, *Genetta genetta*; and fennec fox, *Vulpes zerda*), were found using Google Images and iNaturalist using both the scientific name and the common name for each species as search terms. This approach, taking advantage of the enormous resource of publicly available images, allows access to a much larger data set than in the published scientific literature or than would be possible to obtain *de novo* for this study. Public image-based methods for character state classification have been used previously, such as in a phylogenetic analysis of felid coat patterns (Werdelin and Olsson 1997) and a catalog of iris color variation in the white-browed scrubwren (Cake 2019). However, this approach does require implementing strong criteria for selecting images.

Criteria used to choose images included selecting images where the animal was facing towards the camera, at least one eye was unobstructed, the animal was a non-senescent adult, and the eye was not in direct light (causing glare) or completely in shadow (causing unwanted darkening). The taxonomic identity of the animal in each selected image was verified through images present in the literature, as well as the “research grade” section of iNaturalist. When possible, we collected five images per taxon, although some rarer taxa had fewer than five acceptable images available. In addition, some species with a large number of eye colors needed more than five images to capture their variation, determined by quantitative methods discussed below. Each of the 56 taxa and the number of images used are given in Supplementary Table 2.

Once the images were selected, they were manually edited using MacOS Preview. This editing process involved choosing the “better” of the two eyes for each image (i.e. the one that is most visible and with the least glare and shadow). Then, the section of the iris for that eye without obstruction, such as glare, shadow, or fur, was cropped out. An example of this is given in Figure S11. The strict selection criteria and image editing eliminated the need to color correct the images, a process that can introduce additional subjectivity; the consistency of the data can be seen in the lack of variation between eyes identified as the same color (Figure S5). This process resulted in a data set of 290 cropped, standardized, irises. These images, along with the original photos, can be found in the Supplementary Material.

### Eye color identification

To impartially identify the eye color(s) present in each felid population, the data set images were loaded by species into Python (version 3.8.8) using the Python Imaging Library (PIL) (Van Rossum and Drake 2009; Clark 2015). For each image, the red, green, and blue (RGB) values for each of its pixels were extracted. Then, they were averaged and the associated hex color code for the average R, G, and B values was printed. The color associated with this code was identified using curated and open source color identification programs (Aerne 2022; Cooper 2022). There is no universally agreed upon list of colors, since exact naming conventions differ on an individual and cultural basis, but these programs offer a workable solution, consisting of tens of thousands of colors names derived from published, corporate, and governmental sources. This data allowed the color of each eye in the data set to be impartially assigned, removing a great deal of the bias inherent in a researcher subjectively deciding the color of each iris.

Eye colors were assigned on this basis to one of five fundamental color groups: brown, green (including hazel), yellow (including beige), gray, and blue. The possible color groups were determined before observation of the data based on basic color categories established in the literature: white, black, red, green, yellow, blue, brown, purple, pink, orange, and gray (Berlin and Kay 1991). Of course, not all of the eleven categories ended up being represented by any irises; no irises were observed to be white, black, red, purple, pink, or orange.

As an example of this method, if an iris’s color had the RGB values R: 114, G: 160, B: 193, this would correspond to the hex code #72A0C1. This hex code, when put into the color identification programs, results in the identification “Air Superiority Blue”, derived from the British Royal Air Force’s official flag specifications (Cooper 2022; Aerne 2022). Based on the identification, this iris would be added to the “blue” color group, bypassing a researcher having to choose the color themself. If a color’s name did not already contain one of the eleven aforementioned color categories, the name was searched for in the Inter-Society Color Council-National Bureau of Standards (ISCC–NBS) System of Color Designation (Judd and Kelly 1939). For instance, the color with RGB values R: 37, G: 29, B: 14 corresponds to hex code #251D0E, identified as “Burnt Coffee” by the color identification programs. The ISCC–NBS descriptor for this color is “moderate brown”, so the color would be added to the “brown” group. All colors were able to be placed directly from their color name or their ISCC–NBS descriptor and, for colors with both a color category in the name and an ISCC–NBS descriptor, there were no instances in which the two conflicted.

While color itself lies on a spectrum, splitting the colors into discrete fundamental groups is the most tractable approach to analyze eye color in a biologically reasonable way. If every eye color was instead taken together on one spectrum and analyzed as a continuous trait, the results would be highly unrealistic. As an example, if there were two sister taxa, one with blue eyes (R: 0, G: 0, B: 139) and one with brown eyes (R: 150, G: 75, B: 0), a continuous reconstruction would assign the ancestor the intermediate eye color in the color space: R: 75, G: 37, B: 69.

However, this color is firmly within the “purple” category. It is highly unlikely that a recent ancestor of two taxa with blue and brown eyes had purple eyes, rather than blue eyes, brown eyes, or both, which would be the result if blue and brown were considered as separate categories. Indeed, one would run into the same issue if categories were removed at an earlier stage and each taxon was only considered to have one eye color, determined by averaging all irises. A taxon with blue and brown eyes would again be said to have purple eyes, a color which none of the members of that taxon have. The data being separated into color groups is the most realistic way to investigate this trait, preventing the loss of variation present in the natural populations and simultaneously creating impossible analyses. The lines between color categories are not always clear to an observer (e.g. grayish-blues and bluish-grays can look alike) and, no matter how they are defined, they may still be arbitrary. Nevertheless, this is why we used color identification programs, impartially defining the lines to make the analysis possible.

To ensure no data was missed due to low sample size, the first 500 Google Images, as well as all the “research grade” images on iNaturalist, were manually viewed for each species, while referring back to already analyzed data and periodically checked with the color identification programs (Aerne 2022; Cooper 2022). Any missed colors were added to the data set. This method nonetheless has a small, but non-zero, chance to miss rare eye colors that are present in species. However, overall, it provides a robust and repeatable way to identify the general iris colors present in animals.

In addition, if, for a given species, one or two eye colors were greatly predominant in the available data online (i.e. the first 500 Google Images, as well as all the “research grade” images on iNaturalist), they were defined as being the most common eye color(s). The specific cutoffs were >80% for one eye color or ∼40% for both eye colors respectively. With this assessment, the phylogenetic analysis below could be carried out with all recorded eye colors, as well as using only the most common eye colors, thereby assuring that rare eye colors did not skew the results.

### Color polymorphism assessment

Although placing the eyes in the data set into discrete color groups is useful for downstream analyses, we also wanted to make sure a polymorphic assessment of the iris color trait reflects the reality of the trait. To do this, we performed a principal component analysis (PCA) on the R, G, and B values of every pixel of every iris in the data set, using the package scikit-learn (version 1.2.0) in Python and the built-in stats package in R (version 4.2.1) (Pedregosa *et al*. 2011; R Core Team 2022). The utility of PCA for color polymorphism assessment has been demonstrated before (Paterson and Blouin-Demers 2017). By averaging the pixels, irrespective of color group, in twenty equally spaced bins along each of the first three principal components (PCs), we were able to get a sense of what aspect of the color variation each PC was capturing. Then, we fit a linear mixed model for eye color on each of the PCs using the R package lme4 (version 1.1.34), including the species and individual the pixels were coming from as nested random effects (Bates *et al*. 2015).

This method allowed us to compare the effect of assigned iris color along each principal component axis using Satterthwaite’s t-test with Bonferroni correction. A significant effect of color group for a given PC in the linear mixed model would indicate that the color category assigned according to the methods above is meaningfully predictive of a pixel’s value along that PC. As an example, if the real irises are not adequately represented by the discrete color categories proposed here (e.g. brown eyes are all brownish-gray and gray eyes are all grayish-brown, so the categories significantly overlap), then there should not be a significant effect of assigned eye color for a PC that separates pixels by color (e.g. a PC that separates gray and brown). Of course, due to the nature of converting not fully standardized photographs into pixel data, there are many individual pixels that are outliers within a given eye—for instance, a brown-colored pixel might show up in a eye categorized as “blue” because of a fleck of dust in the cat’s eye or some irregular pigmentation—but, unless these outliers are numerous (thus making them not outliers), they should not affect this analysis.

If a significant effect of a color group was found for a PC, the PC values for all categories were compared to one another using a post-hoc Tukey HSD test with Bonferroni correction from the package emmeans (version 1.8.8), in order to distinguish which groups in particular significantly differ for that PC (Lenth 2023). Although this analysis was able to determine which color groups adequately reflect the true trait distributions and are meaningful overall, this does not necessarily mean that a polymorphic view of eye color is appropriate for all species. To address this, since PC2 was demonstrated to be the axis that separates pixels by relevant colors, the pixels in each color group for each species were compared along PC2 using a Kruskal–Wallis test with Bonferroni correction to determine whether there was a significant effect of color group at all. If there was, a two-sided pairwise Mann-Whitney-Wilcoxon test with Bonferroni correction was used to compare each group to one another. In this way, we were able to determine the biological appropriateness of using discrete color categories to analyze felid iris color.

### Shade measurements within each color group

Although averaging the pixels within each iris was sufficient to categorize the colors present for each felid taxon, not every felid iris has homogeneous pigmentation. For example, some colors in some taxa are subject to central heterochromia with a darker pigment near the pupil and a lighter pigment in the periphery (Figure 1b, h) or the reverse (Figure 1c). Thus, we calculated corresponding “shade” values for each color group in each species. To do this, the images were sorted into their color groups for each species. For each group, RGB values for each pixel in each image were again extracted, resulting in a three dimensional data set. This was reduced to two dimensions using Uniform Manifold Approximation and Projection (UMAP), a method selected for its preservation of local structure, important for potential fine shade differences (McInnes *et al*. 2018). The UMAP projection for each image was then analyzed using k-means clustering through scikit-learn (Pedregosa *et al*. 2011). The number of clusters (k), indicating the number of distinct shades of color in the iris of each animal, was determined using elbow plots.

After this was done for all images in the group, the k values were averaged and each image was clustered using the average k value, rounded to the nearest integer. This was done to standardize within groups, avoid confounders based on lower quality images, and allow for comparative analysis. After this, the average RGB values for each cluster for each image were calculated. Then, the clusters were matched up based on similarity. To do this, one image from the group had its clusters labeled in order (if there were three clusters, they would be 0, 1 and 2). Then, another image from the group would have the distances in 3D space between each of its clusters compared to each of the labeled clusters. The optimal arrangement of clusters was found by calculating the sum of squared errors for every possible combination of clusters and taking the minimum. Then, the clusters were merged. This method was repeated for every image in the group. Doing this for every color of every species resulted in an output with the number of shades within the iris for each color in each species, as well as an average of each different shade across the data. Throughout this process, images were not resized so as to allow higher quality images with more pixels to contribute a greater amount to the average. This was done to ensure any blurring from lower quality images did not obscure the true shade variety in each eye.

The final, combined clusters were ranked by how prevalent they were within the eyes, calculated by the number of pixels in each group, and the groups for each shade were categorized as “dark”, “medium”, or “light”. To do this, if there were three general clusters for a color of a species, the distance from black (RGB: 0,0,0) in 3D space for each of the cluster average RGB values was computed and then they were assigned to be “dark”, “medium”, or “light” based on increasing distance from black in the color space. For species with two shades in their eyes of a certain color, the cluster average RGB values were compared, again using distance, to the averages of the three-shade eye “dark”, “medium”, and “light” values. They would be assigned the label that they were closest to. The remaining space was filled: if “dark” or “light” was empty, the “medium” value was duplicated; if “medium” was empty, the “dark” and “light” values were averaged. This method allows two-shade eyes to be compared to three-shade eyes without losing vital information. For species with one shade of a color in their eyes (of which there ended up being none in the data set), its average RGB values were assigned to “dark”, “medium”, and “light”. Lastly, eyes with four shades had to combine the two most similar shades together in order to make them comparable to the rest of the data set. The importance of this pipeline is to create a data set that can be compared in a standardized way. The information about which shades are most represented was also collected and saved. This data can be found in the Supplementary Material.

To ensure these results were accurately assessing eye color, the RGB values for each shade within each species were compared with increasing numbers of images from the data set (for examples, see Figure S5). If the RGB values leveled off as sample size increased, that would indicate that the sample is representative of the “true” shades. If there were major fluctuations, that would indicate that the sample size is not high enough to overcome variations in lighting conditions. In this way, the sample sizes for each color present in each taxon were confirmed to be sufficient as their RGB values leveled off.

### Phylogeny

The phylogeny used for this work was a subset from the Carnivora supertree from Nyakatura and Bininda-Emonds (2012). This ultrametric phylogeny takes into account 188 literature and gene trees and includes members of all eight Felidae lineages. More recent phylogenies are largely congruent, differing mainly in the placement of the Bay Cat Lineage and the Pallas’s cat (*Otocolobus manul*), partly due to differences in Y chromosome evolutionary evidence compared to other lines of evidence (Li *et al*. 2016). Alternate placements were tested and were found to not produce a significant difference in results, making these discrepancies irrelevant to this study.

This Carnivora supertree tree is missing nine of the extant felid groups for which data was collected. Thus, a second tree (termed the “full” tree) was created with the missing species being added manually according to their placements on a Felidae specific tree from Johnson *et al*. (2006) and/or the more recent tree from Li *et al*. (2016). The subspecies added were defined according to the most recent identification based on Kitchener *et al*. (2017) and Liu *et al*. (2018). Subspecies were added as a polytomy next to the previously defined species on the tree. Since divergence data was unavailable for some of the species and subspecies, the additions were made with branch lengths equal to the nearest resolved neighboring branch, a severe overestimation of the divergence between groups.

It is important to note that this method of manually adding taxa to a tree is flawed without proper sequence data and certainly should not be relied upon for ancestral state predictions or to make broad claims, as there is no guarantee that any addition reflects true divergence. However, this tree was created purely to provide some insight into local areas of the tree at the species level (e.g. what was the eye color of the ancestral tiger?). Even still, these predictions must be understood as far more uncertain than analyses with the original supertree with more limited taxa. The full tree with all the eye colors present for each species is shown in Figure 2. The main tree created only considering the most common eye colors is presented in Figure S3a-e.

### General color reconstruction

To begin the process of ancestral state reconstruction, the phylogenetic trees were read into R using the package ape (version 5.6-2) (Paradis and Schliep 2019). A table of taxa, and the colors represented for each, was loaded in and scored with 0/1 for absence/presence. The same table with just the most common eye colors was also loaded in.

The optimal model of trait evolution was then determined for each of the five eye colors independently across the tree. This was done using an Akaike information criterion (AIC) analysis done on the results of 14 different models run using the R package phytools (version 1.2-0) (Revell 2012). Six of the models were run using equal/symmetric rates (ER): a continuous-time Markov (Mk) model using fitMK(); an Mk model with edge rates assumed to have been sampled randomly from a Γ distribution using fitgammaMk(); a hidden rates model with four hidden states using fitHRM(); a hidden rates model with six hidden states using fitHRM(); a hidden rates model with a possible hidden state when a color is present, but not when it is absent, using fitHRM(); and a hidden rates model with a possible hidden state when a color is absent, but not when it is present, using fitHRM(). Another six of the models were identical, except they were run using asymmetric rates (ARD). The final two used an Mk model, but assumed that a color cannot be lost after it is gained or that it cannot be gained after it is lost, respectively. This process was done for the data of all the observed eye colors, as well as for the data for the most common eye colors and for the full phylogeny, with the AIC output and weights for the ER and ARD models given in Supplementary Table 3. The model with the lowest AIC value, indicating the best explanation of the data given the number of parameters, was used for subsequent analyses. The best models were also rerun in corHMM (version 2.8) and no differences between the results were found (Beaulieu *et al*. 2022).

Although the presence/absence of each eye color were analyzed on their own, the colors are likely not fully independent. Therefore, they were also analyzed together as a polymorphic trait using stochastic mapping through fitpolyMk() in phytools. Since there were far too many states (2^5^-1), including high parameter complexity, for adequate interpretation as a polymorphic character and the two analyses generally aligned (data not shown), the independent model was used. A color was said to be present at any given node (Figure 3) if the marginal maximum likelihood ancestral state reconstruction for that color was greater than 50%, indicating more support for presence than absence.

### Quantitative color reconstruction

After data was collected on the eye colors present for every node on the tree, more specific reconstructions were possible. For each node, a new tree was created for each eye color present at that node. Each of these subset trees included every descendant of that node that shared each eye color with it, except for those where the color was lost and then re-arose independently. For example, an ancestral node that was determined to have green eyes and brown eyes present would have one tree with all its continuous, green-eyed descendants and another tree with all its continuous, brown-eyed descendants. A diagram of this method is given in Figure S12. This method was done to most accurately reconstruct along plausible evolutionary pathways. If one wants to predict the eye shade of a specific color for a specific node, one should omit taxa that either have lost that eye color (since their present condition cannot communicate any relevant information about the shade of that color for their ancestor), as well as taxa that have lost that eye color and then regained it (since it is unknown whether their present condition is at all related to the shade of that color for their ancestor).

After the trees were created, the specific colors were reconstructed using maximum likelihood methods with the function fastAnc() from phytools (Revell 2012). This was done independently for the red, green, and blue values for each of the data sets collected for the light, medium, and dark shades. Since RGB values can only be from 0-255, it was heartening that the 95% confidence intervals for the quantitative reconstructions were almost always well within the realistic range, lending considerable support to the reconstructions. Large confidence intervals are a known limitation of continuous trait likelihood reconstructions, so one should not understand the reconstructions to always communicate the exact eye shades of the felid ancestors, but they are useful in comparison to one another to illuminate larger trends.

Beyond reconstructing the colors themselves, corHMM’s rayDISC() was used to reconstruct the number of shades within each eye color for each node, using the shade representation data as a discrete, multistate trait (Beaulieu *et al*. 2022). This was also done for the primary and secondary shades within each eye. Put together, these methods allow for a high resolution understanding of the iris color of ancestral felids. For each ancestral felid population, we are able to know: which color eyes were present (out of brown, green, yellow, gray, and blue), how many different shades they had in their eyes for each color, which shades were more or less common, and approximately what those shades would have been.

### Environmental/behavioral/physical trait encoding

Data on pupil shape was obtained from Banks *et al*. (2015) and data on activity by time of day and primary habitat(s) was obtained from the University of Michigan Animal Diversity Web (Banks *et al*. 2015; Myers *et al*. 2022). Data on zoogeographical regions were based on Johnson *et al*. (2006) and data on coat patterns were based on Werdelin *et al*. (1997). Nose color data (pink or black) and whether or not any black was present in the coat or tail were determined manually from observation of images.

For correlation comparisons, each multistate trait was converted into a set of binary traits. Pupil shape was scored with 0 for vertical/subcircular pupils and 1 for round pupils. Likewise, whether black coloration is present in the coat and whether black coloration is present in the tail were scored with a 0 for absence and a 1 for presence. Activity was split into three traits, each corresponding to an activity lifestyle: nocturnal, crepuscular, and diurnal. Then, for each felid taxon, each trait was scored as present or absent using 1 or 0, respectively. This was especially useful, given that some taxa fall into multiple categories. The same was done for historical zoogeographical region (nearctic [North America], neotropical [South America], palearctic [Europe and North Asia], oriental [South Asia], ethiopian [Africa]), primary habitat (mountains, rainforest, forest, savanna, desert), and coat pattern (flecks, uniform, stripes, rosettes, blotches, sblotch [small blotches]). This method was also used for pink and black nose colors because some felid noses contain both colors and some species have both of the colors represented in their populations; in both cases, both colors would be marked as present.

### Correlation analysis

Apart from reconstructing ancestral states, different correlations were performed in order to investigate the possible evolutionary interactions related to eye color variation. The environmental/physical trait data, along with the presence/absence data for each eye color, was analyzed with a maximum likelihood approach using BayesTraits (version 3.0.5), made accessible in R through the package btw (version 2.0) (Pagel *et al*. 2004; Griffin 2018). This was done by building two models, one where the evolution of two binary traits is independent and one where their evolution is dependent on one another (i.e. where the rate of change in one trait is influenced by the state of the other trait). Then, the models were evaluated using a calculated log Bayes Factor, with a log Bayes Factor over 2 indicating positive evidence for the dependent model. Given the stochasticity of these models, the model comparisons were done 100 times and the calculated log Bayes Factors were averaged, ensuring robust and reproducible results. This process was done by comparing the presence of each eye color to all others, as well as the environmental/behavioral/physical data to the presence of each eye color, the average shade of the RGB values in each eye color, and the average shade of the RGB values in all eye colors overall. This latter average was computed for all taxa by dropping NA values in the averages. To transform the average values into discrete traits, each value was categorized using Jenks natural breaks optimization, performed through the getJenksBreaks() command in the package BAMMtools (version 2.1.10) (Rabosky *et al*. 2014). Finally, tetrachoric correlation coefficients were calculated using the tetrachoric() command in the package psych (version 2.2.9), to indicate the direction of each association (Revelle 2022). For the shade correlations, a positive association indicates that the trait is associated with lighter shades.

## Literature Cited

1. Aerne D. 2022. Color Names. GitHub repository; [accessed 2022 Nov 18] https://github.com/meodai/color-names

2. Banks MS, Sprague WW, Schmoll J, Parnell JAQ, Love GD. 2015. Why do animal eyes have pupils of different shapes? Sci. Adv. 1(7):e1500391.

3. Bates D, Maechler M, Bolker B, Walker S. 2015. Fitting Linear Mixed-Effects Models Using lme4. J. Stat. Softw. 67(1): 1–48.

4. Berlin, B, Kay P. Basic color terms: Their universality and evolution. Univ of California Press, 1991.

5. Cake M. 2019. Regional variation in iris colour of the White-browed Scrubwren’Sericornis frontalis’ complex in digital photographs. Aust. Field Ornithol. 36: 148–153.

6. Clark A. 2015. Pillow (PIL Fork) Documentation. readthedocs. [accessed 2023 Feb 10] https://buildmedia.readthedocs.org/media/pdf/pillow/latest/pillow.pdf

7. Clark DL, Clark RA. 2016. Neutral point testing of color vision in the domestic cat. Exp. Eye Res. 153:23–26.

8. Cooper R. 2022. Color Namer. GitHub repository; [accessed 2022 Nov 18] https://github.com/robertcoopercode/color-namer

9. Corbett EC, Brumfield RT, and Faircloth BC. 2023. The mechanistic, genetic and evolutionary causes of bird eye colour variation. Ibis 10.1111/ibi.13276

10. Cote J, Le Galliard JF, Rossi JM, Fitze PS. 2008. Environmentally induced changes in carotenoid-based coloration of female lizards: a comment on Vercken et al. J. Evol. Biol. 21(4): 1165–72.

11. Craig AJFK, and Hulley PE. 2004. Iris colour in passerine birds: why be bright-eyed? S. Afr. J. Sci. 100(11): 584–588.

12. Davison A, Jackson HJ, Murphy EW, Reader T. 2019.Discrete or indiscrete? Redefining the colour polymorphism of the land snail Cepaea nemoralis. Hered. 123(2): 162–75.

13. Donnelly MP, Paschou P, Grigorenko E, Gurwitz D, Barta C, Lu RB, Zhukova OV, Kim JJ, Siniscalco M, New M, and Li H. 2012. A global view of the OCA2-HERC2 region and pigmentation. Hum. Genet. 131: 683–696.

14. Draus-Barini J, Walsh S, Pośpiech E, Kupiec T, Głąb H, Branicki W, Kayser M. 2013. Bona fide colour: DNA prediction of human eye and hair colour from ancient and contemporary skeletal remains. Investig. Genet. 4: 1–5.

15. Frost P. 2006. European hair and eye color: a case of frequency-dependent sexual selection?. Evol. Hum. Behav. 27(2):85–103.

16. Griffin RH. 2018. _btw: Run BayesTraitsV3 from R_. R package version 2.0.

17. iNaturalist. [accessed 2022 Nov 20] https://www.inaturalist.org.

18. Ito S, Wakamatsu K, Ozeki H. 2000. Chemical analysis of melanins and its application to the study of the regulation of melanogenesis. Pigment Cell Res. 13: 103–9.

19. Jablonski NG, and Chaplin G. 2017. The colours of humanity: the evolution of pigmentation in the human lineage. Philos. Trans. R. Soc. 372(1724): 20160349.

20. Johnson WE., Eizirik E, Pecon-Slattery J, Murphy WJ, Antunes A, Teeling E, O’Brien SJ. 2006. The late Miocene radiation of modern Felidae: a genetic assessment. Science 311(5757):73–77.

21. Judd DB., and Kelly KL. 1939. Method of designating colors and a dictionary. J. Res. Natl. Inst. Stand. Technol. 23:355–385.

22. Kolb H, Fernandez E, Nelson R. 2011. Webvision: the organization of the retina and visual system [Internet]. Salt Lake City (UT): University of Utah Health Sciences Center.

23. Lenth R. 2023. emmeans: Estimated Marginal Means, aka Least-Squares Means. https://CRAN.R-project.org/package=emmeans, Version = 1.8.8.

24. Li G, Davis BW, Eizirik E, Murphy WJ. 2016. Phylogenomic evidence for ancient hybridization in the genomes of living cats (Felidae). Genome Res. 26(1):1–11.

25. Liu, F, Wollstein, A, Hysi, PG, Ankra-Badu, GA, Spector, TD, Park, D, Zhu, G, Larsson, M, Duffy, DL, Montgomery, GW, Mackey, DA. 2010. Digital quantification of human eye color highlights genetic association of three new loci. PLoS Genet. 6(5): e1000934.

26. Liu Y-C, Sun X, Driscoll C, Miquelle DG, Xu X, Martelli P, Uphyrkina O, Smith JDL, O’Brien SJ, Luo S-J. 2018. Genome-wide evolutionary analysis of natural history and adaptation in the world’s tigers. Current Biology 28(23): 3840–3849.

27. Lyons LA. 2012. Genetic testing in domestic cats. Mol. Cell. Probes. 26(6):224–230.

28. McInnes L, Healy J, Melville J. 2018. Umap: Uniform manifold approximation and projection for dimension reduction. arXiv preprint. arXiv:1802.03426.

29. Merola M. 1994. A reassessment of homozygosity and the case for inbreeding depression in the cheetah, Acinonyx jubatus: implications for conservation. Conserv. Biol. 8(4): 961–71.

30. Mould MC, Huet M, Senegas L, Milá B, Thébaud C, Bourgeois Y, Chaine AS. 2023. Beyond morphs: Inter-individual colour variation despite strong genetic determinism of colour morphs in a wild bird. J. Evol. Biol. 36(1): 82–94.

31. Myers P, Espinosa R, Parr CS, Jones T, Hammond GS, Dewey TA. 2022. The Animal Diversity Web (online). [accessed 2022 Dec 1]. https://animaldiversity.org.

32. Negro JJ, Carmen Blázquez M, Galván I. 2017. Intraspecific eye color variability in birds and mammals: a recent evolutionary event exclusive to humans and domestic animals. Front. Zool. 14(1):1–6

33. Nyakatura, K, Bininda-Emonds ORP. 2012. Updating the evolutionary history of Carnivora (Mammalia): a new species-level supertree complete with divergence time estimates. BMC Biology 10: 1–31.

34. Oliveira R, Godinho R, Randi E, Ferrand N, Alves PC. 2008. Molecular analysis of hybridisation between wild and domestic cats (Felis silvestris) in Portugal: implications for conservation. Conserv. Genet. 9(1):1–11.

35. Pagel M, Meade A, Barker D. 2004. Bayesian Estimation of Ancestral Character States on Phylogenies. Syst. Biol. 53(5):673–684.

36. Paradis E, Schliep K. 2018. ape 5.0: an environment for modern phylogenetics and evolutionary analyses in R. Bioinformatics. 35(3):526–528.

37. Passarotto A, Parejo D, Cruz-Miralles A, Avilés JM. 2018. The evolution of iris colour in relation to nocturnality in owls. J. Avian Biol. 49(12).

38. Paterson JE, Blouin-Demers G. 2017. Distinguishing discrete polymorphism from continuous variation in throat colour of tree lizards, Urosaurus ornatus. Biol. J. Linn. Soc. 121(1): 72–81.

39. Pedregosa F, Varoquaux G, Gramfort A, Michel V, Thirion B, Grisel O, Blondel M, Prettenhofer P, Weiss R, Dubourg V, Vanderplas J. 2011. Scikit-learn: Machine learning in Python. J. Mach. Learn. Res. 12(Oct):2825–2830.

40. R Core Team. 2022. R: A language and environment for statistical computing. R Foundation for Statistical Computing, Vienna, Austria.

41. Rabosky DL, Grundler M, Anderson C, Title P, Shi JJ, Brown JW, Huang H, Larson JG. 2014. BAMMtools: an R package for the analysis of evolutionary dynamics on phylogenetic trees. Methods Ecol. Evol. 5(7):701–707.

42. Revell LJ. 2012. phytools: an R package for phylogenetic comparative biology (and other things). Methods Ecol. Evol. 3(2):217–223.

43. Revelle W. 2022. psych: Procedures for Personality and Psychological Research, Northwestern University, Evanston, Illinois, USA, https://CRAN.R-project.org/package=psych, Version = 2.2.9.

44. Simcoe M, Valdes A, Liu F, Furlotte NA, Evans DM, Hemani G, Ring SM, Smith GD, Duffy DL, Zhu G, and Gordon SD. 2021. Genome-wide association study in almost 195,000 individuals identifies 50 previously unidentified genetic loci for eye color. Sci. Adv., 7(11): p.eabd1239.

45. Strain GM. 2007. Deafness in blue-eyed white cats: The uphill road to solving polygenic disorders. J. Vet. Med. 173(3):471–472.

46. Van Rossum G, Drake FL. 2009. Python 3 Reference Manual. Scotts Valley, CA: CreateSpace.

47. Vercken E, Sinervo B, Clobert J. 2008. Colour variation in female common lizards: why we should speak of morphs, a reply to Cote et al. J. Evol. Biol. 21(4): 1160–4.

48. Werdelin L, Olsson L. 1997. How the leopard got its spots: a phylogenetic view of the evolution of felid coat patterns. Biol. J. Linn. Soc. 62(3):383–400.

49. White D, Rabago-Smith M. 2010. Genotype–phenotype associations and human eye color. J. Hum. Genet. 56(1):5–7.

50. Wilhelmy J, Serpell J, Brown D, Siracusa C. 2016. Behavioral associations with breed, coat type, and eye color in single-breed cats. J. Vet. Behav. 13:80–87.

