## Supplemental Results for "Evolutionary Insights Into Felidae Iris Color Through Ancestral State Reconstruction"

*Model fits:* In the main analysis, the best models for trait evolution were found to be: an All Rates Different extended Mk model for brown and blue eyes, meaning that the transition from absence to presence and presence to absence had different rates; an Equal Rates Mk model for green eyes, meaning that the transition rates from absence to presence and presence to absence were identical; an Equal Rates model with a hidden rate when the color is absent for yellow eyes; and an All Rates Different model with a hidden rate when the color is absent for gray eyes (Supplementary Table 3).

*Quantitative reconstruction data spread:* For each color, the average standard deviation of the resulting RGB shade values over the data set was calculated. This measure of the spread of colors present over the species considered was about the same for brown, green, and yellow eyes, with values between 22.1 and 26.9. It is remarkable that the interspecies variability in eye color is approximately the same for these three different colors. However, the average standard deviation was 31.6 and 31.2 for gray and blue eyes, respectively, demonstrating substantially more variability, although the latter can be explained by the low number of taxa with blue eyes. In much the same way, calculating the average standard deviation for light, medium, and dark shades across the data set resulted in values of 26.9, 21.6, and 17.2 respectively. There is increasing variability across species as shades get lighter. When all red, green, and blue values are taken separately across the data set, the average standard deviations are 42.3, 40.5, and 39.2 respectively. Thus, the three basic colors that encode the more complex colors in the data set vary between species to the same degree, an important validation of the method used in this paper, given that it is not at all obvious that RGB values would encode biologically relevant data.

Since R, G, and B values vary to the same degree, there is no evidence of asymmetricality in the importance of each factor for encoding the eye colors.

*Yellow, green, and blue shade reconstructions:* The evolution of yellow eye shades has fewer defined patterns, due to its more recent evolution (Figure S6). The yellow that emerged in the ancestor of the *Panthera* genus was a brownish yellow, as was the yellow that emerged in the ancestor of the Lynx, Leopard Cat, Domestic, and Puma Lineages. In extant species with the yellow eye color, the brown content in this color is sometimes increased (such as for *Puma concolor*) and sometimes greatly decreased (such as for *Lynx lynx*). For green and blue eye shades, there is even less evidence of a pattern, given the lack of ancestral nodes with the predicted presence of those colors (Figures S7 and S8). An interesting aspect of all three of these colors is that there are a number of species that developed multiple different colors in one eye, rather than (or in addition to) multiple shades. This likely accounts for the majority of the outlier/overlapping points in Figures S1 and S2. An example of this is the blue of *Panthera uncia* (snow leopards). Snow leopard blue eyes routinely have a thin outer band of beige and an inner layer of blue. Our program was easily able to pick this up in its dimensionality reduction, but it should be noted that the reconstruction shades do not have the same level of contrast that the original eyes might have had. This is a result of averaging samples and could be corrected through significantly more data. This issue does not occur for *Panthera pardus*, whose eyes have a similar makeup with a beige outer ring and an inner blue (Figure 1h). The program is able to output that the beige area is generally about the same size for *Panthera pardus* (primary shade) than for *Panthera uncia* (also primary shade). This method of bypassing what otherwise would have been painstaking morphological measurements is another benefit of this methodology and program.

*Correlation analysis supplement:* There were numerous significant correlations for zoogeographical regions. Three correlations stand out for their strength: brown eyes-nearctic (BF = 3.77, corr = 0.97), gray eyes-palearctic (BF = 8.39, corr = 0.99), and blue eyes-neotropical in the two alternate analyses (BF = 4.02/4.70, corr = -0.95/-0.96). Every nearctic felid has brown eyes, every palearctic felid has gray eyes, and no neotropical felid has blue eyes. These are not the only such uniform associations on the tree, but they are the ones unable to be explained by evolutionary history and phylogeny. They are also not easily directly explained by the various habitats present in the regions, given the sparse number of significant associations between eye colors and habitats. The only trend of note is that blue eyes are strongly associated with forests (BF = 2.43, corr = 0.96), with almost all of the species with blue eyes occupying this habitat to some degree.

The traits relating to body pigmentation (coat pattern, black fur present on the tail and the body, and nose color) also revealed few correlations with eye color presence. Having a pink nose is strongly negatively correlated with brown eyes (BF = 6.32, corr = -0.98), the eye color with the highest amount of eumelanin.[1] Likewise, the eye color with the lowest amount of melanin (blue) is strongly negatively correlated with having a black nose (BF = 4.05, corr = -0.98). Having a coat pattern with blotches or small blotches is highly related to ancestry and these traits are not evenly distributed across the phylogeny.[2] Because of this, the correlation coefficients between these traits and various eye colors are usually very large (near 1 or -1) and almost entirely explainable by phylogeny, due to coat pattern often being a lineage specific trait.

---

[1] Kolb H, Fernandez E, Nelson R. 2011. Webvision: the organization of the retina and visual system [Internet]. Salt Lake City (UT): University of Utah Health Sciences Center.

[2] Werdelin, Lars, and Lennart Olsson. "How the leopard got its spots: a phylogenetic view of the evolution of felid coat patterns." *Biological Journal of the Linnean Society* 62, no. 3 (1997): 383-400.

However, two such correlations cannot purely be explained by phylogeny: brown-small blotches (BF = 2.54, corr = 0.97) and green-blotches (BF = 2.32, corr = -0.97).

Correlation analysis was also performed for the physical/behavioral/environmental data and the continuous average RGB shade values. This was done because some factors could affect the shades within eye colors, regardless of whether or not they affect the presence of eye colors categories as a whole (Figure S10). Many factors are only significantly correlated with one aspect within certain eye colors (out of red, green, and blue shades). Additionally, there does not seem to be much consistency for the effect of various factors across eye colors. For instance, living in the palearctic zoogeographical region is associated with darker reds and greens in brown eyes (BF = 2.86, corr = -0.98; BF = 2.27, corr = -0.98), but lighter reds and greens in yellow eyes (BF = 6.31, corr = 1; BF = 7.84, corr = 1). Despite discordances like this, with all eye colors taken together, the palearctic region is highly correlated with lighter shades of red, green, and blue (BF = 4.63, corr = 0.55; BF = 7.66, corr = 0.99; BF = 2.83, corr = 0.46). This type of complexity is found in most of the factors considered in this study.

Nevertheless, broad trends can be observed. First, the RGB values of green eyes showed the greatest sensitivity to environmental/behavioral/physical factors (20 significant correlations), followed by yellow (19), brown (17), blue (14), then gray (11). For brown eyes, the shades of red and green, the two colors that make up RGB brown, are what are most often affected by the environmental/behavioral/physical factors, and always in the same direction and magnitude, an expected result and a good check of the method. For green, the green shades are understandably the most affected by the environmental/behavioral/physical factors. These results provide additional confidence that the representation of natural colors in a computerized RGB encoding retains biologically relevant information.

The overall shades of red, green, and blue across all eye colors are affected in similar ways. The presence of brown eyes is significantly correlated with darker overall shades for all RGB axes for a species. On the other hand, the presence of yellow eyes is correlated with lighter overall red and green shades. The presence of blue or gray eyes is correlated with lighter overall green and blue shades. Living in a palearctic (and often, colder) environment is significantly associated with overall lightening of eyes in all RGB aspects. Of course, these associations only indicate evidence for correlated evolution; the direction of the correlation is often less clear.
