## Supplemental Figures for "Evolutionary Insights Into Felidae Iris Color Through Ancestral State Reconstruction"

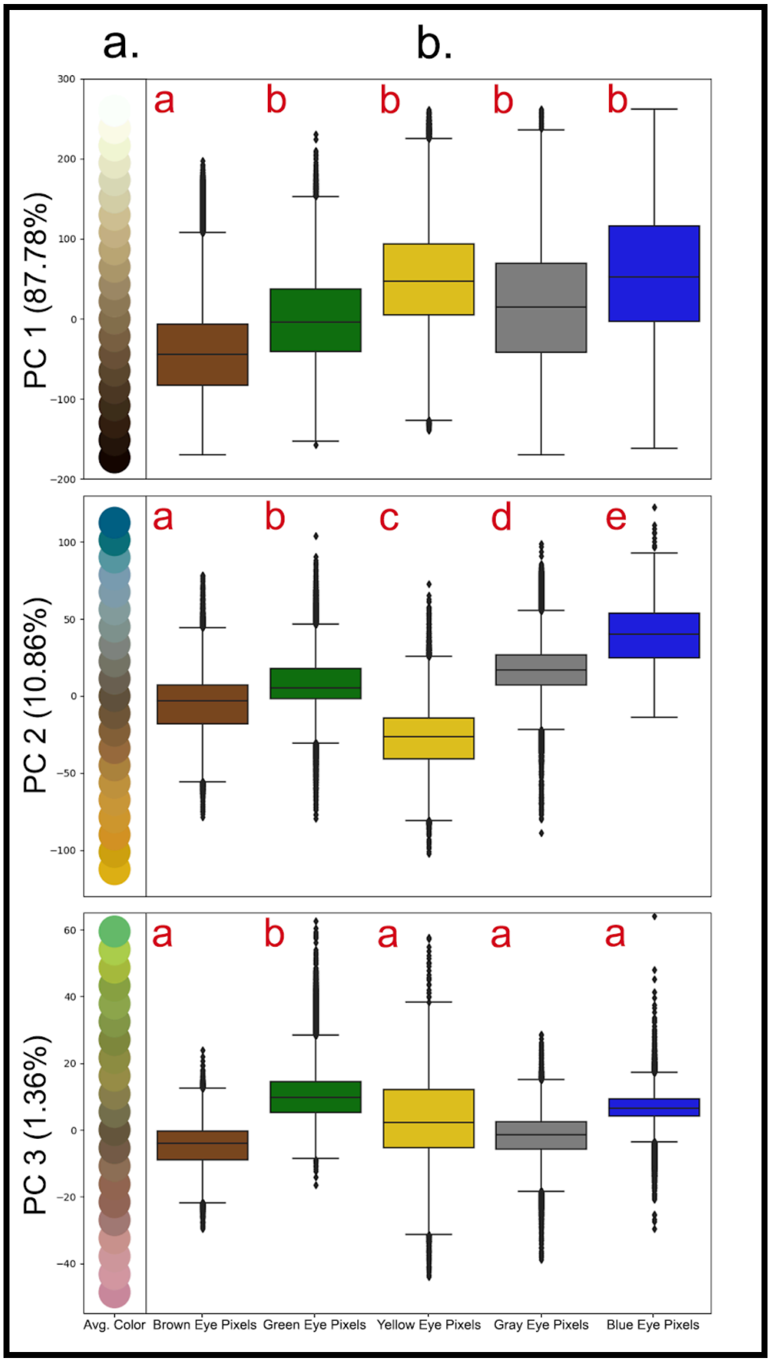


Figure S1: (a) The average pixel color along PCs 1-3. PCA performed for the RGB values for all pixels in the data set. Color average presented was calculated for the same data in 20 equally spaced bins along each PC. (b) Distribution of pixels from all eyes in the data set for each of the color categories for each PC. Different letters indicate groups with significant differences ($p < 0.05$) according to post-hoc Tukey HSD test, after comparing linear mixed models with Satterthwaite's t-test, both tests with Bonferroni correction.

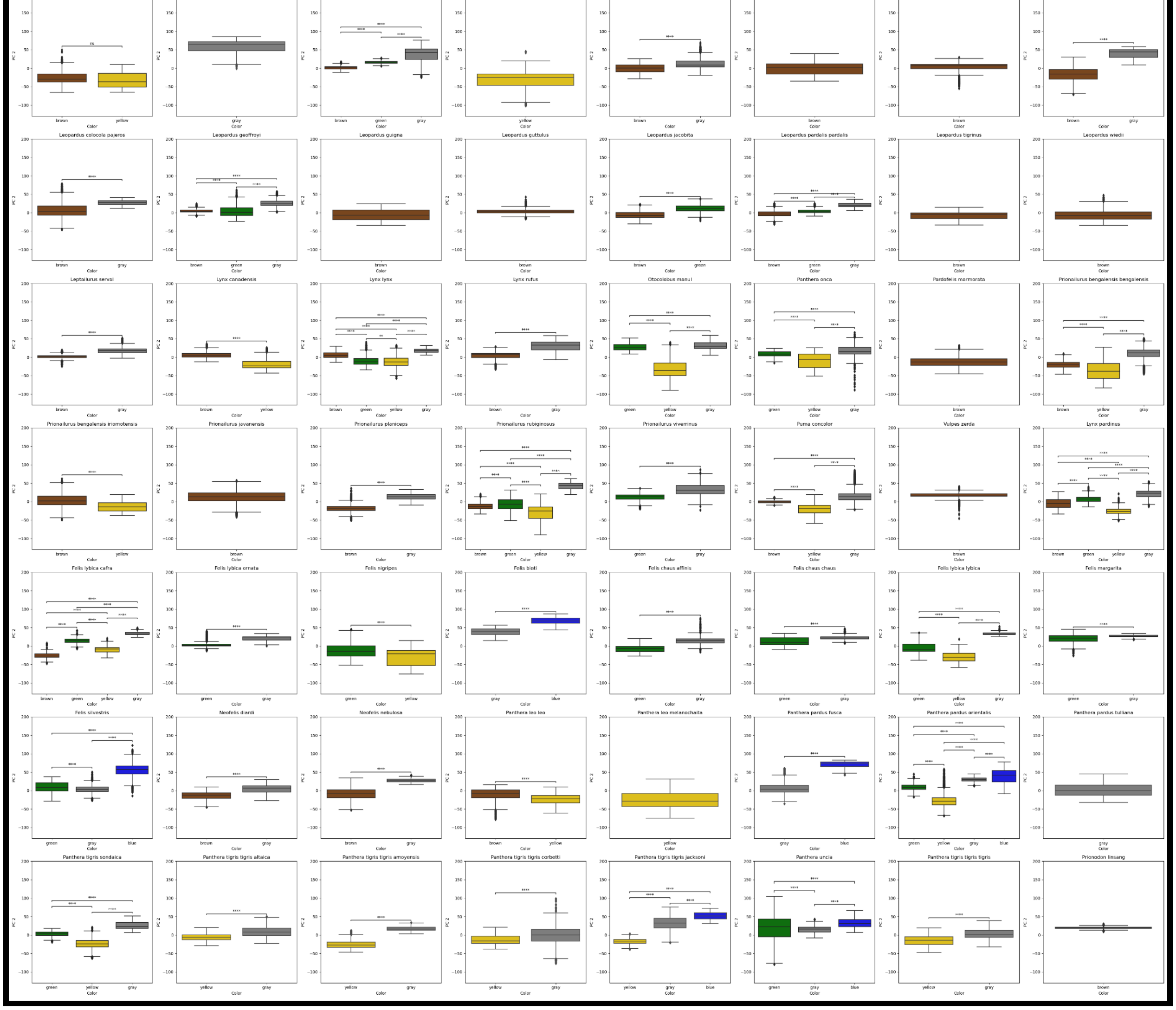

Figure S2: Distribution of pixels for each species for each of the color categories for PC 2. Significances calculated using pairwise Mann-Whitney-Wilcoxon tests, two-sided with Bonferroni correction. ns: $p > 0.05$; *: $0.05 > p > 0.01$; **: $0.01 > p > 0.001$; ***: $0.001 > p > 0.0001$; ****: $0.0001 > p$.

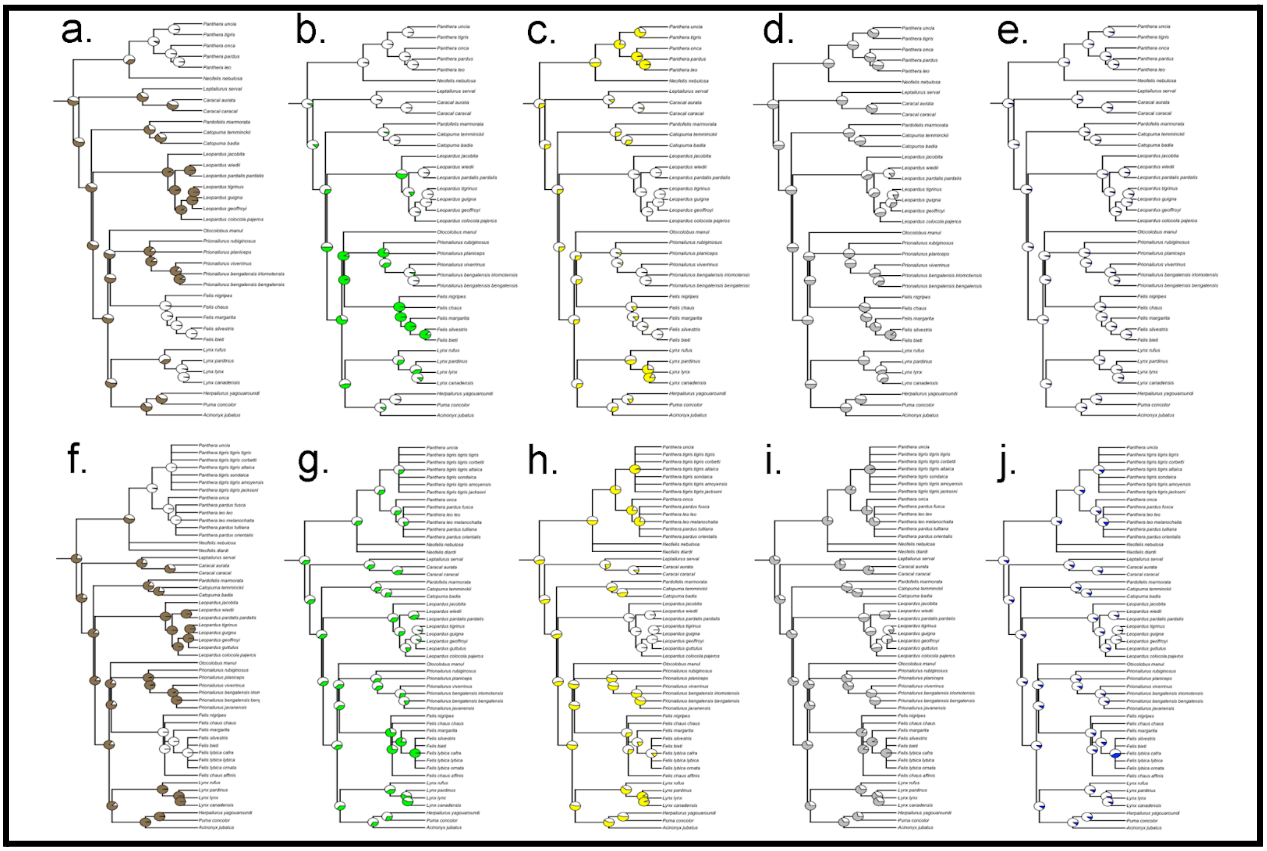


Figure S3: Maximum likelihood probabilities for the ancestral states of brown, green, yellow, gray, and blue eye colors when only the most common eye colors were considered (a-e) or all of the subspecies were added to the tree (f-j). The amount of color in the node pie charts represents the support for that color being present at that node. Exact branch lengths are not plotted.

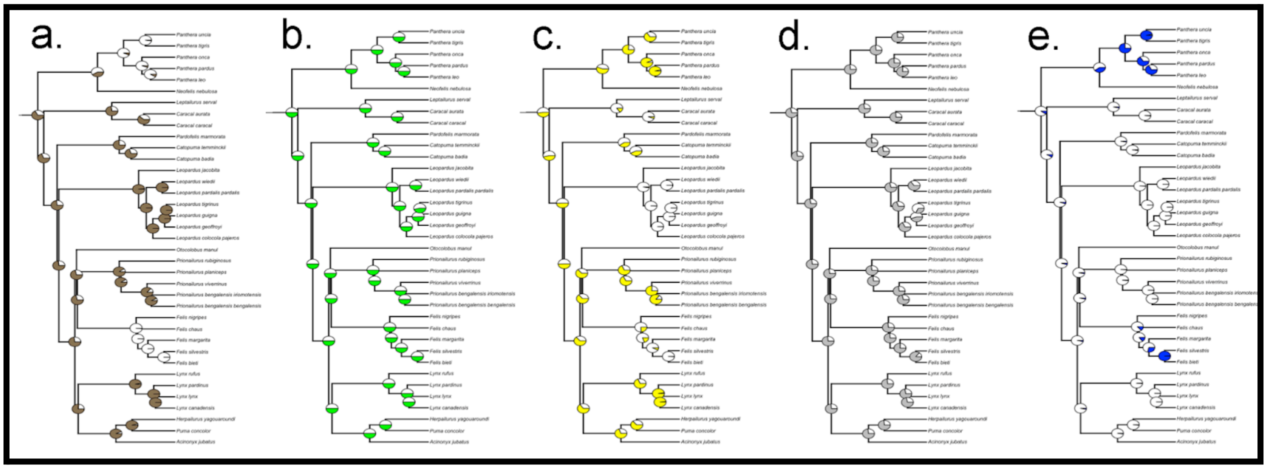


Figure S4: Maximum likelihood probabilities for the ancestral states of brown (a), green (b), yellow (c), gray (d), and blue (e) eye colors. The amount of color in the node pie charts represents the support for that color being present at that node. Exact branch lengths are not plotted.

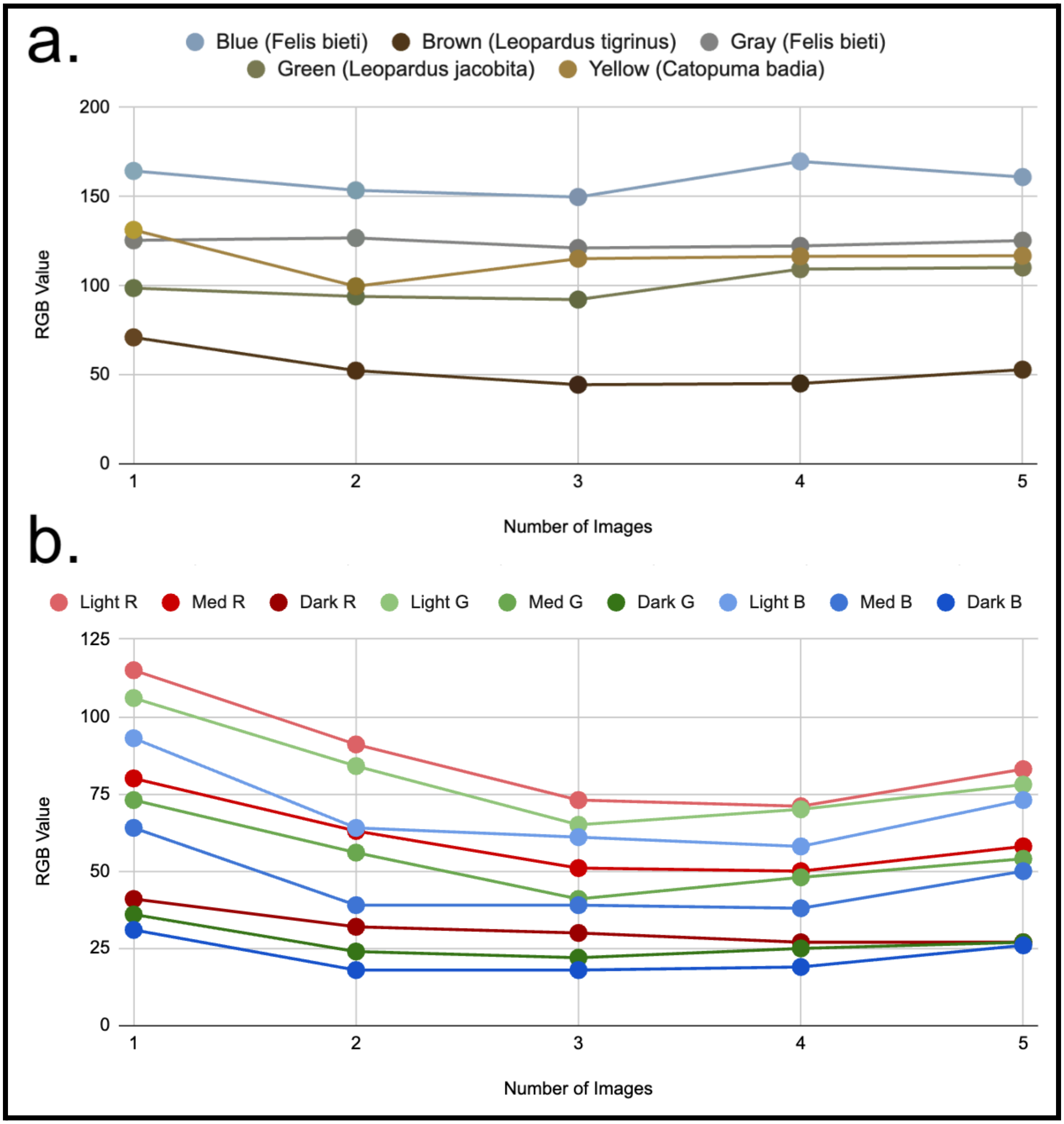


Figure S5: (a) Average RGB values by the number of images used for representative image sets: all brown-eyed individuals from *Leopardus tigrinus*, all green-eyed individuals from *Leopardus jacobita*, all yellow-eyed individuals from *Catopuma badia*, and all gray- and blue-eyed individuals from *Felis bieti*. RGB values displayed were averages between the R, G, and B values for all shades. Point colors correspond to the average eye color for that number of images. (b) RGB values by the number of images used for the dark, medium, and light shades present in the brown eyes of *Leopardus tigrinus*. The leveling off in both parts of the figure indicates that the sample is sufficient.

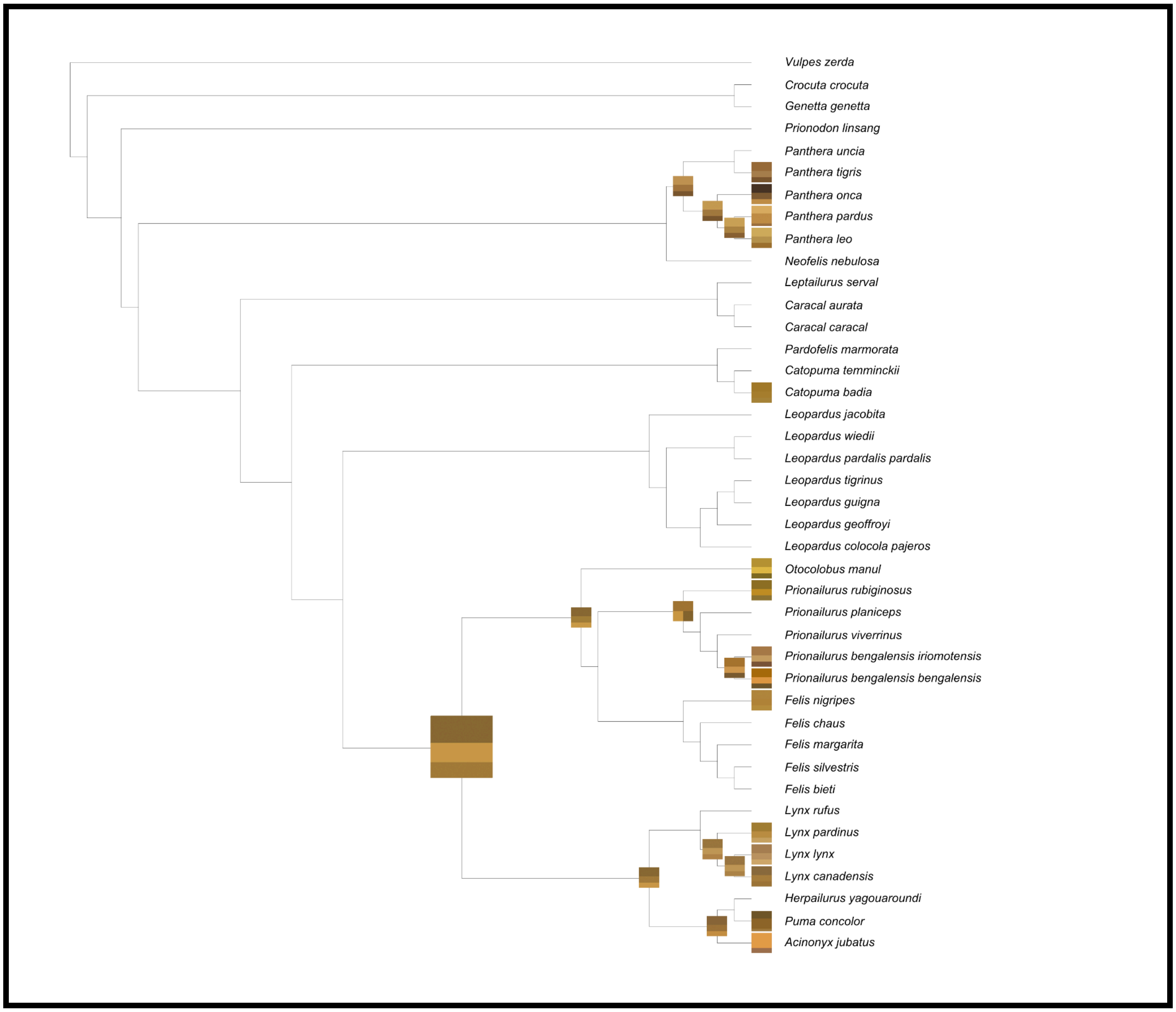


Figure S6: Reconstruction of the ancestral states of the shades of yellow eyes. The squares at each node are the quantitative reconstructed shades. The proportion of a square that a shade takes up indicates how common that shade is in the data. Exact branch lengths are not plotted.

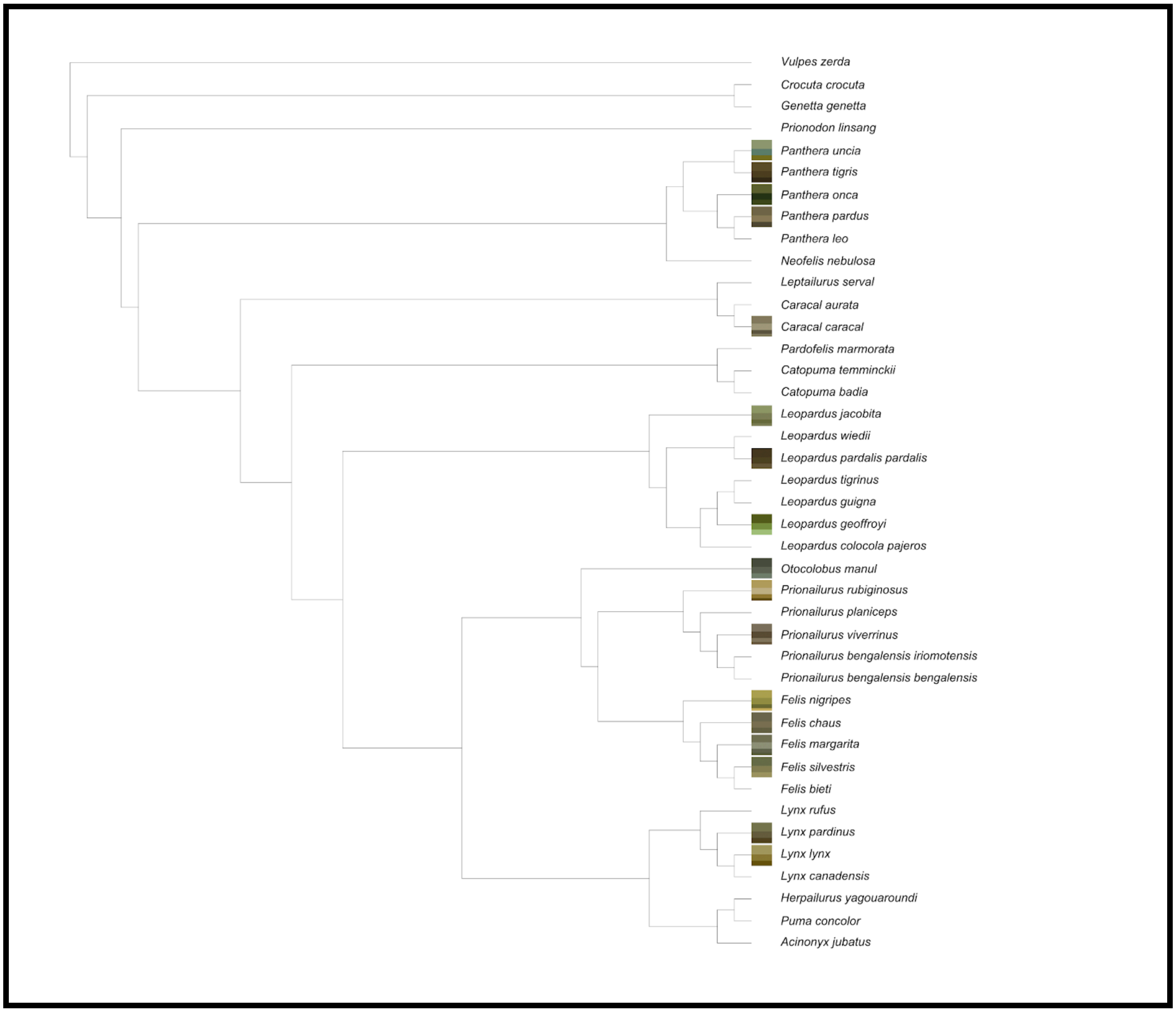


Figure S7: Reconstruction of the ancestral states of the shades of green eyes. The squares at each node are the quantitative reconstructed shades. The proportion of a square that a shade takes up indicates how common that shade is in the data. Exact branch lengths are not plotted.

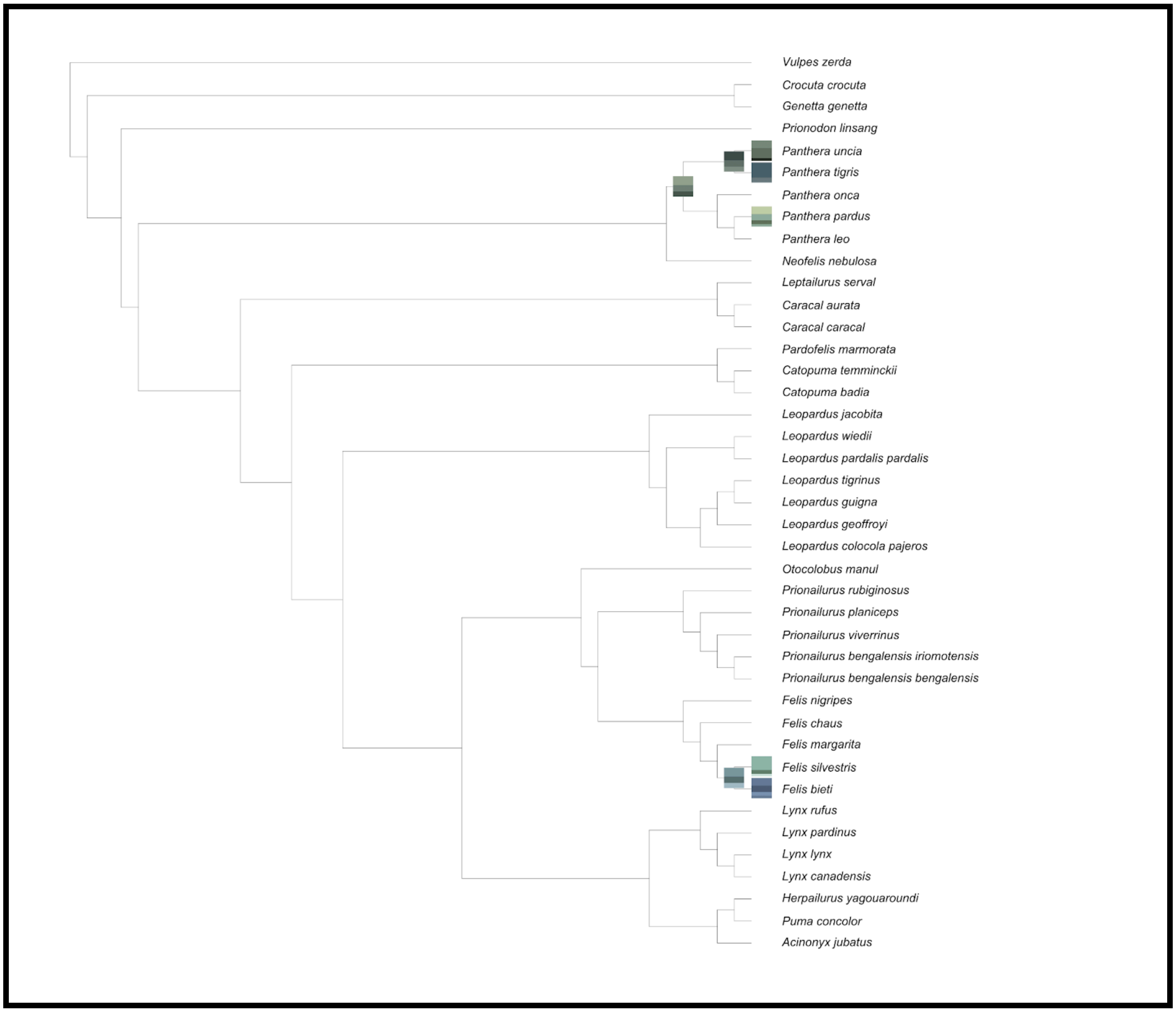


Figure S8: Reconstruction of the ancestral states of the shades of blue eyes. The squares at each node are the quantitative reconstructed shades. The proportion of a square that a shade takes up indicates how common that shade is in the data. Exact branch lengths are not plotted.

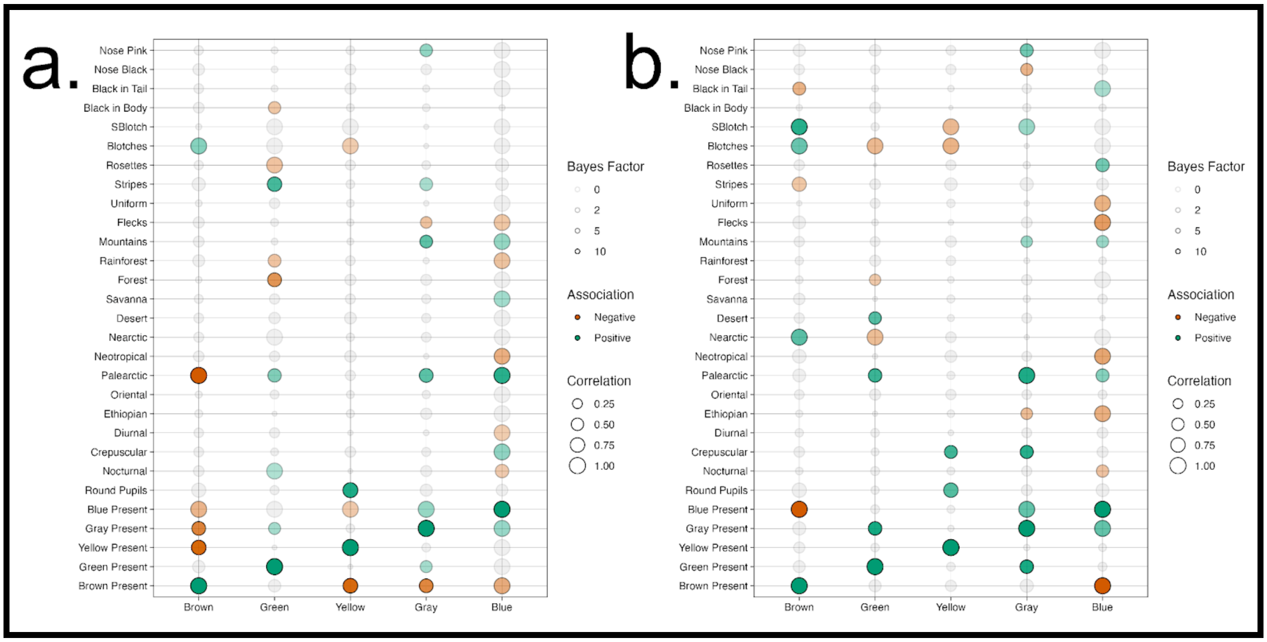


Figure S9: Correlations between the presence of each eye color and various physical, behavioral, and environmental factors for just the most common eye colors (a) and when all of the subspecies were added to the tree (b). Larger circles correspond to stronger correlations and more opaque circles correspond to more significant correlations. Green circles have a positive correlation, red circles have a negative correlation, and gray circles do not meet the significance threshold (Bayes factor = 2).

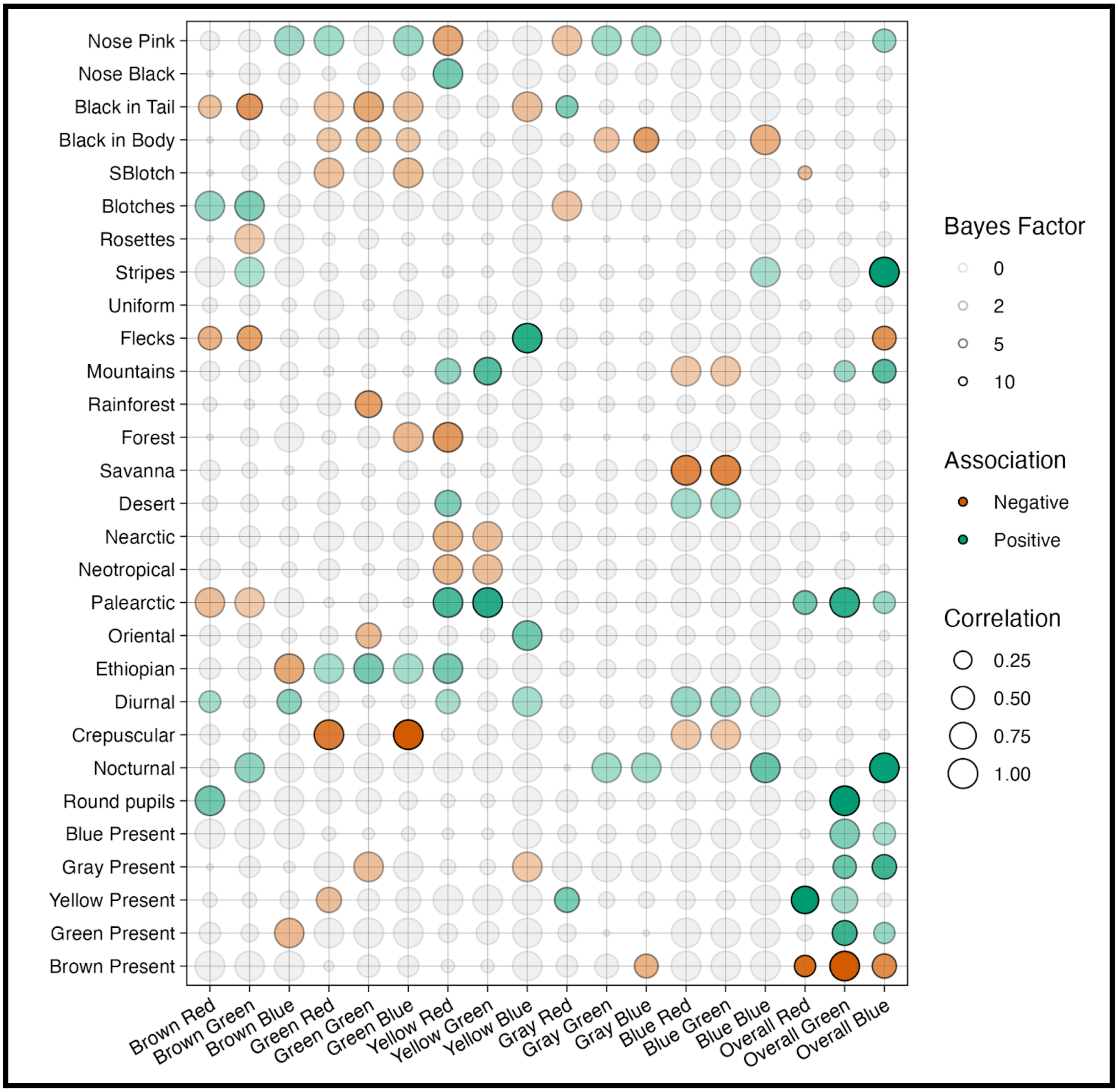


Figure S10: Correlations between the shades of the RGB values of each eye color and various physical, behavioral, and environmental factors. Larger circles correspond to stronger correlations and more opaque circles correspond to more significant correlations. Green circles have a positive correlation (i.e. lighter shade), red circles have a negative correlation (i.e. darker shade), and gray circles do not meet the significance threshold (Bayes factor = 2).

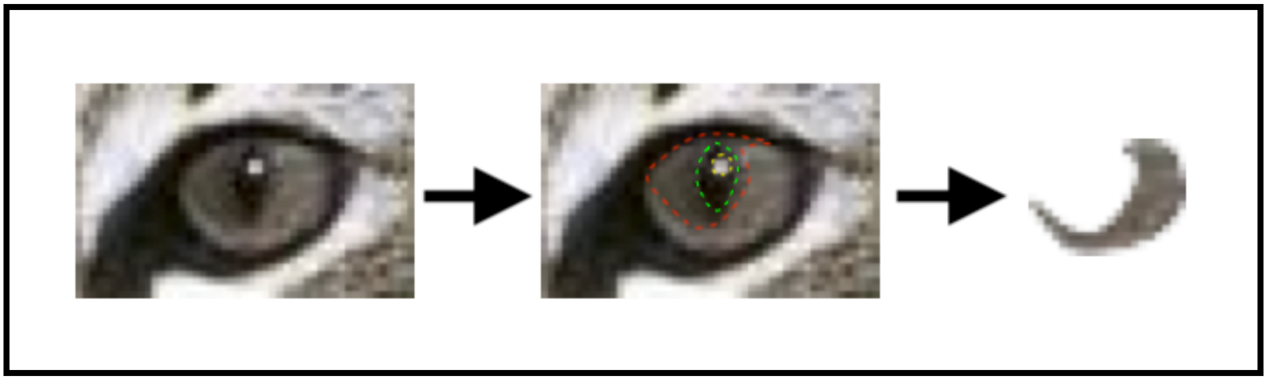

Figure S11: Example of the iris cropping process. On the left is an eye of *Leopardus geoffroyi*, the Geoffroy's cat. In the middle are dotted lines around parts of the image to cut out: red - shadow, green - pupil, yellow - glare. On the right is the resulting iris used for analysis. Credit to dreamstime.com for the original photo.

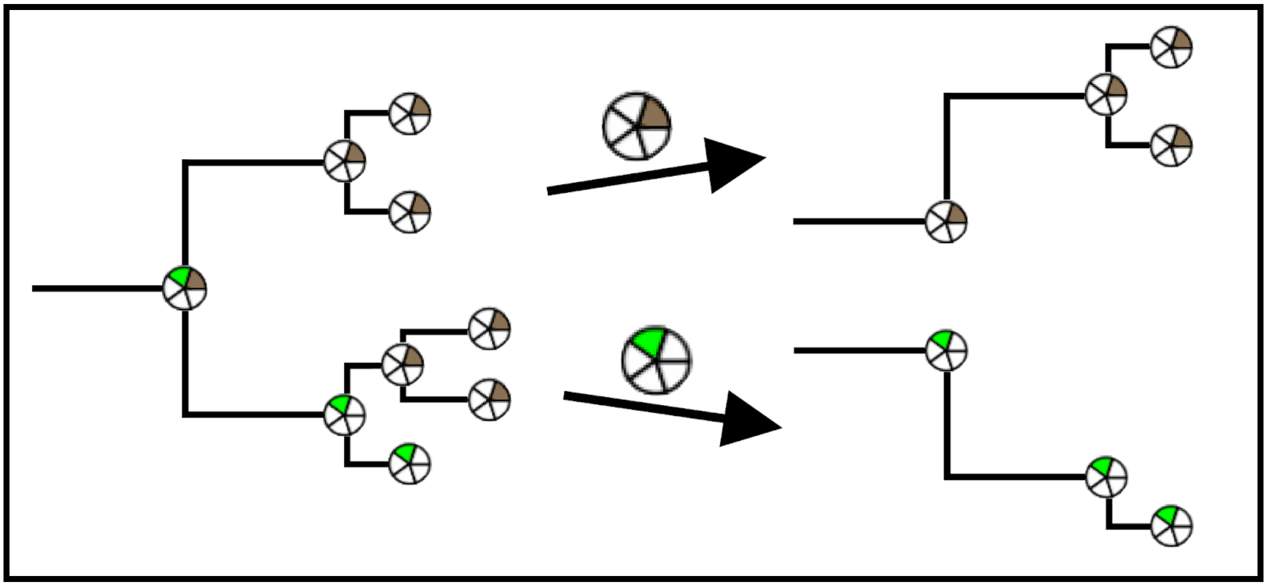

Figure S12: An example phylogenetic tree with illustration of tree separation for shade analysis. The five-wedge pie charts indicate presence (color) or absence (white) of various iris colors. Here, the ancestral node has brown and green eyes. The reconstruction for that node's brown eye shades, shown after the upper arrow, include all the continuous, brown-eyed descendants. The green reconstruction, after the bottom arrow, is done the same way.
